# The long noncoding RNA Charme supervises cardiomyocyte maturation by controlling cell differentiation programs in the developing heart

**DOI:** 10.1101/2022.07.05.498800

**Authors:** Valeria Taliani, Giulia Buonaiuto, Fabio Desideri, Adriano Setti, Tiziana Santini, Silvia Galfrè, Leonardo Schirone, Davide Mariani, Giacomo Frati, Valentina Valenti, Sebastiano Sciarretta, Emerald Perlas, Carmine Nicoletti, Antonio Musarò, Monica Ballarino

**Author notes:** Corresponding author: Monica Ballarino e-mail address. These authors contributed equally to this work (Valeria Taliani, Giulia Buonaiuto).

## Abstract

Long noncoding RNAs (lncRNAs) are emerging as critical regulators of heart physiology and disease, although the studies unveiling their modes-of-action are still limited to few examples. We recently identified pCharme, a chromatin-associated lncRNA whose functional knockout in mice results in defective myogenesis and morphological remodelling of the cardiac muscle. Here, we combined Cap-Analysis of Gene Expression (CAGE), single-cell (sc)RNA sequencing and whole-mount *in situ* hybridization analyses to study pCharme cardiac expression. Since the early steps of cardiomyogenesis, we found the lncRNA being specifically restricted to cardiomyocytes, where it assists the formation of specific nuclear condensates containing MATR3, as well as important RNAs for cardiac development. In line with the functional significance of these activities, pCharme ablation in mice results in a delayed maturation of cardiomyocytes, which ultimately leads to morphological alterations of the myocardium and ventricular hypo-trabeculation. Since congenital anomalies in myocardium are clinically relevant in humans and predispose patients to major complications, the identification of novel genes controlling cardiac morphology becomes crucial. Our study offers unique insights into a novel lncRNA-mediated regulatory mechanism promoting cardiomyocyte maturation and bears relevance to Charme locus for future theranostic applications.

## INTRODUCTION

In all vertebrates, heart development occurs through a cascade of events in which a subtle equilibrium between proliferation, migration, and differentiation ultimately leads precursor cells to mature into all the major cardiac cell types (Bruneau 2013; Moorman and Christoffels 2003). At the molecular level, the execution of the developmental program is governed by the dynamic interplay of several cardiac regulators, whose expression and functions are coordinated in time and space. Alteration of this process results in abnormal cardiac morphogenesis and other congenital heart defects that in humans represent the most common types of birth defects and cause of infant death (Zimmerman et al., 2020; Srivastava, 2006; Center for Disease Control and Prevention, 2020). Mutations of transcription factors and their protein cofactors have emerged as causative of a broad spectrum of heart malformations, although only 15-20% of all congenital heart defects have been associated to known genetic conditions (Morton et al., 2022; Kodo et al., 2009; Bouveret et al., 2015; Ang et al., 2016). The need to find more targets for diagnosis and therapy has therefore evoked interest in new categories of disease-causing genes, whose discovery has been accelerated by impressive advancements in genomics. In this direction, improvements in the next-generation sequencing technologies have revolutionised many areas of the biomedical research, including cardiology, by the discovery of long noncoding RNA (lncRNAs) (Mattick et al., 2004; Cipriano et al., 2018, Rinn and Chang 2012; Ulitsky and Bartel 2013; Yao et al., 2019). LncRNAs form a heterogeneous class of non-protein coding transcripts, longer than 200 nucleotides, participating in many physiological (i.e. cell stemness, differentiation or tissue development) and pathological (i.e. cancer, inflammation, cardiovascular or neurodegeneration) processes (Azad et al., 2021; Hu et al., 2018; Riva et al., 2016). In the heart, several biologically relevant lncRNAs have been identified, with functions related to aging (Jusic et al., 2022) and regeneration (Pagano et al., 2020; Wang et al., 2021). Specifically for cardiac development (Pinheiro et al., 2021), most of the lncRNA-mediated mechanisms were dissected by cell culture studies (Klattenhoff et al., 2013; Ounzain et al., 2015; Kim et al., 2021), while *in vivo* characterizations remained limited to few instances (Grote et al. 2013; Anderson et al., 2016; Ritter et al., 2019; Hazra,R,et al., 2022).

The importance of lncRNA for adult heart homeostasis has been often described by linking their aberrant expression to cardiac anomalies (Han et al., 2014; Wang et al., 2016; Ballarino et al., 2018; Han et al., 2019; Ponnusamy et al., 2019; Anderson et al., 2021). Along this line, in 2015 we identified a new collection of muscle-specific lncRNAs (Ballarino et al., 2015) and focused our attention on Charme (Chromatin architect of muscle expression) (mm9 chr7:51,728,798-51,740,770), a mammalian conserved lncRNA gene, whose loss-of-function in mice causes progressive myopathy and, intriguingly, congenital heart remodelling (Ballarino et al., 2018). Here, we look at the function of this lncRNA in a developmental window and assign an embryonal origin to the cardiac defects produced by its knockout. By using molecular and whole-tissue imaging approaches, we give an integrated view of Charme locus activation in the developing heart and show that the functional pCharme isoform, retaining intron-1, is expressed already in the fetal tissues, progressively dropping down after birth. Single-cell (sc)RNA sequencing analyses from available datasets also reveal a strong cell type-specific expression of pCharme in the embryonal heart, where it is restricted to cardiomyocytes (CM). In line with this specific location, high-throughput transcriptomic analyses combined with the phenotypic characterization of WT and Charme^KO^ developing hearts, highlight a functional role of the lncRNA in CM maturation and in the developmental formation of trabeculated myocardium. Furthermore, high-throughput sequencing of RNA isolated from fetal cardiac biopsies upon cross-linking and immunoprecipitation (CLIP) of MATR3, a nuclear pCharme interactor (Desideri et al., 2020), reveals the existence of a specific RNA-rich condensate containing pCharme as well as RNAs for important regulators of embryo development, cardiac function, and morphogenesis. In accordance with the functional importance of the pCharme/MATR3 interaction in developing cardiomyocytes, the binding of these transcripts with MATR3 was altered in Charme^KO^ hearts. Moreover, MATR3 depletion in mouse-derived cardiac primary cells leads to the down-regulation of such RNAs, which suggests an active crosstalk between MATR3, pCharme and cardiac regulatory pathways.

Understanding the basics of cardiac development from a lncRNA *point-of-view* is of interest not only for unravelling novel RNA-mediated circuitries but also for improving treatment options aimed at enhancing cardiomyogenic differentiation. Indeed, despite the recent advancements in generating cardiomyocytes from pluripotent stem cells through tissue engineering-based methods, most protocols produce immature cells, which lack many attributes of adult cardiomyocytes (Uosaki et al., 2015). Consequently, the cells generated cannot be used for efficient drug screening, modelling of adult-onset disease, or as a source for cell replacement therapy. We identify here pCharme as a new non-coding regulator of cardiac maturation and characterise the molecular interactome acting with the lncRNA *in vivo*. This research not only advances our understanding of heart physiology but in the next few years may serve as a foundation for new human diagnostic and therapeutic approaches.

## RESULTS

### Charme locus expression in developing mouse embryos and cardiomyocytes

Our earlier studies revealed the occurrence of muscle hyperplasia and cardiac remodelling upon the CRISPR/Cas9-mediated Charme loss-of-function in mice (Ballarino et al., 2018; Desideri et al., 2020). Intriguingly, the morphological malformations were clearly displayed in both adult and neonatal mice, strongly suggesting possible roles for the lncRNA during embryogenesis. With the purpose to trace back the developmental origins of Charme functions, we started by analysing the whole collection of FANTOM5 mouse promoterome data to quantify transcription initiation events captured by CAGE-seq across the lncRNA locus (Noguchi et al., 2017). In addition to its cardiac specificity, this profiling revealed the highest expression of the locus during development (E11-E15) followed by a gradual decrease at postnatal stages (**Figure 1A** and **Supplementary File 1**). We then evaluated the expression of the two splice variants, pCharme and mCharme, produced by the locus at E15.5 and postnatally (day 2, after birth). RT-qPCR analyses with specific primers confirmed the CAGE output and revealed that both the isoforms are more expressed at the fetal than the postnatal stage (**Figure 1B**). More importantly, we found pCharme to be 50% more abundant than mCharme, which is particularly intriguing in consideration of the prominent role of the nuclear isoform previously shown in skeletal myogenesis (Desideri et al., 2020). Using *in-situ* hybridization (ISH) approaches, we then profiled by imaging the spatio-temporal expression of Charme during development. These analyses revealed an initial expression of Charme locus already in the tubular heart (**Figure 1C**, E8.5), within territories which will give rise to the future atria and ventricles (V), with the inflow (IFT) and outflow (OFT) tracts displaying the highest signals. ISH also confirmed the expression of the locus at later stages of development, both in cardiac tissues and somites (S) (**Figure 1C**, E13.5; **Figure 1-figure supplement 1A**). A similar expression pattern was found within intact embryos and fetal hearts by whole-mount *in-situ* hybridization (WISH) (**Figure 1D**). To note, specific signals were detected only in wild-type (Charme^WT^) heart muscles, whereas no signal was found in Charme knockout organs (Charme^KO^) (**Figure 1D**, left; **Figure 1-figure supplement 1B**), taken from our previously generated mice (Ballarino et al., 2018). As distinct cell subpopulations are known to form the heart, each one carrying out a specialised function in cardiac development and physiology (de Soysa et al., 2019), we then processed publicly available single-cell RNA sequencing (scRNA-seq) datasets from embryonal hearts (E12.5, Jackson-Weaver et al., 2020) for studying at a deeper resolution the locus expression across individual cell types. Upon clustering (**Figure 1E**, upper) the different cardiac cell-types based on the expression of representative marker genes (Li et al., 2016; Jackson-Weaver et al., 2020; Franco et al., 2006; Meilhac et al., 2018) (**Figure 1-figure supplement 1C-1E**), we found pCharme expression restricted to cardiomyocytes, with an almost identical distribution between the atrial (CM-A), ventricular (CM-V) and other (CM-VP, CM-IVS, CM-OFT) cell clusters (**Figure 1E**, lower).

**Figure 1.**
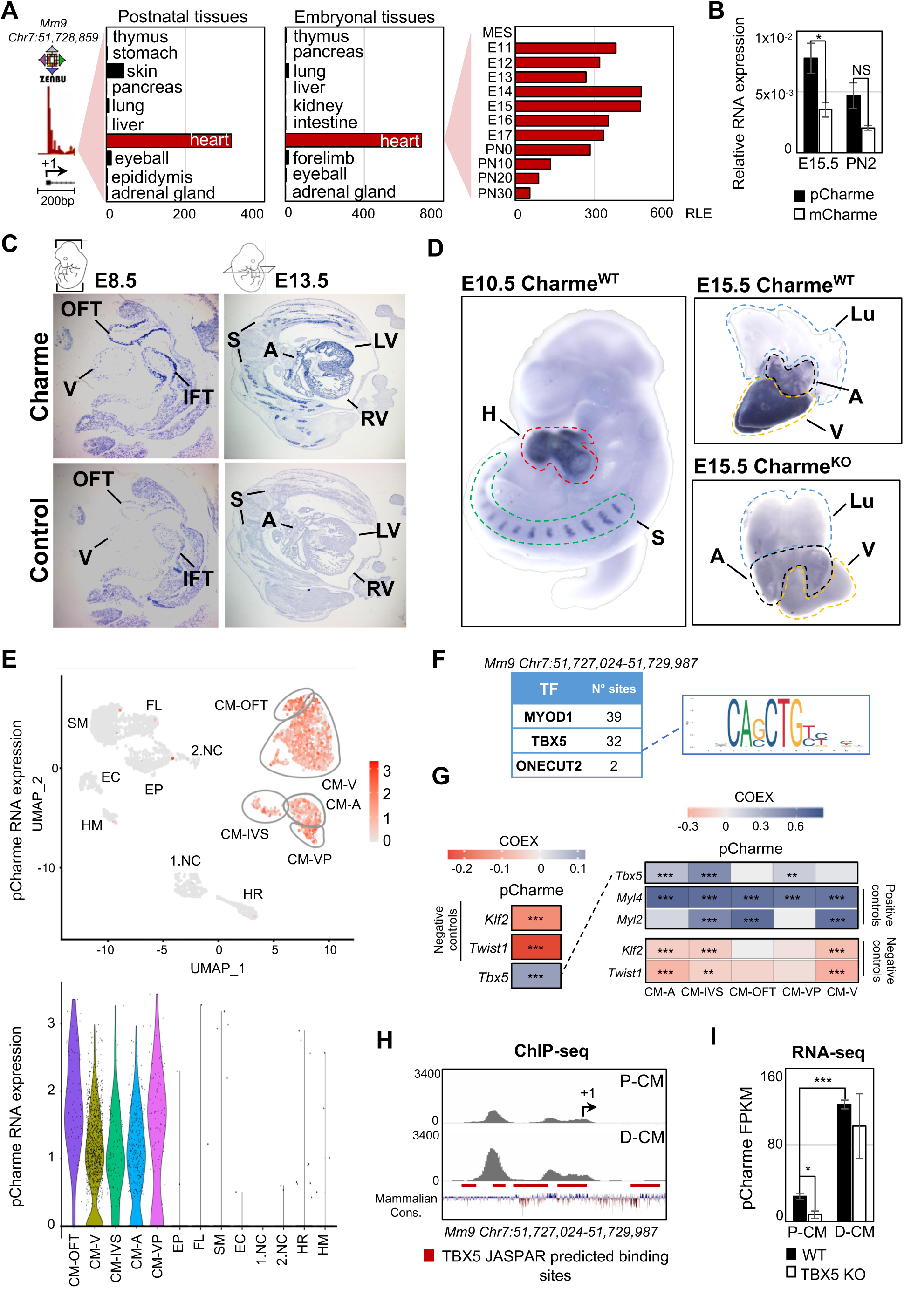
Charme locus expression in developing mouse embryos and cardiomyocytes. **A)** Transcriptional start site (TSS) usage analysis from FANTOM5 CAGE Phase1 and 2 datasets (skeletal muscle is not included) on the last update of Zenbu browser (https://fantom.gsc.riken.jp/zenbu/; FANTOM5 Mouse mm9 promoterome) showing Charme locus expression in postnatal and embryonal body districts (left and middle panels) and during different stages of cardiac development (right panel, E11-PN30). MES=Mesoderm. Bars represent the Relative Logaritmic Expression (RLE) of the Tag Per Million values of Charme TSS usage in the specific samples. **B)** Quantitative Reverse Transcription PCR (RT-qPCR) amplification of pCharme and mCharme isoforms in RNA extracts from Charme^WT^ E15.5 and neonatal (PN2) hearts. Data were normalized to *Gapdh* mRNA and represent means ± SEM of 3 pools. **C)** *In-situ* hybridization (ISH) performed on embryonal cryo-sections using digoxigenin-labelled RNA antisense (Charme, upper panel) or sense (control, lower panel) probes against Charme. Representative images from two stages of embryonal development (E8.5 and E13.5) are shown. OFT: Outflow Tract; IFT: Inflow Tract; V: Ventricle; LV/RV: Left/Right Ventricle; A: Atria; S: Somites. **D)** Whole-mount *in-situ* hybridization (WISH) performed on Charme*^WT^* intact embryos (E10.5, left panel) and Charme*^WT^* and Charme*^KO^* hearts at their definitive morphologies (E15.5, right panels). Signal is specifically detected in heart (H, red line) and somites (S, green line). Lungs (Lu, blue line) show no signal for Charme. The specificity of the staining can be appreciated by the complete absence of signal in explanted hearts from Charme^KO^ mice (Ballarino et al., 2018). A: Atria (black line); V: Ventricles (yellow line). **E)** Upper panel: UMAP plot showing pCharme expression in single-cell transcriptomes of embryonal (E12.5) hearts (Jackson-Weaver et al., 2020). Lower panel: Violin plot of pCharme expression in the different clusters (see Materials and Methods for details). CM: Cardiomyocytes, CM-A: Atrial-CM, CM-V: Ventricular-CM, ISV: Interventricular Septum, VP: Venous Pole, OFT: Outflow Tract, NC: Neural Crest cells, EP: Epicardial cells, FL: Fibroblasts like cells, EC: Endothelial Cells, SM: Smooth Muscle cells, HM: Hemopoietic Myeloid cells, HR: Hemopoietic Red blood cells. **F)** *In silico* analysis of MYOD1, TBX5 and ONECUT2 transcription factors (TF) binding sites using Jaspar 2022 (relative profile score threshold=80%) (Castro-Mondragon et al., 2022). MyoD1 and Onecut2 were used as positive and negative controls, respectively. N° sites =number of consensus motifs. **G)** COTAN heatmap obtained using the whole scRNA-seq (left) and contrasted subsetted dataset (right) showing pCharme and *Tbx5* expression correlation (COEX). *Myl4* and *Myl2* were used as positive controls for cardiomyocytes while *Klf2* and *Twist1* as negative controls (markers of EC and NC, respectively). See Materials and Methods for details. **H)** TBX5 ChIP-seq analysis across Charme promoter in murine precursors (P-CM) and differentiated cardiomyocytes (D-CM) (GSE72223, Luna-Zurita et al., 2016). The genomic coordinates of the promoter, the Charme TSS (+1, black arrow), the TBX5 JASPAR predicted binding sites (red lines) and the mammalian conservation track (Mammalian Cons.) from UCSC genome browser are reported. **I)** Quantification of pCharme expression from RNA-seq analysis performed in wild type (WT) and TBX5 knockout (KO) murine P-CM and D-CM (SRP062699, Luna-Zurita et al., 2016). FPKM: Fragments Per Kilobase of transcript per Million mapped reads. Data information: *p < 0.05; ***p < 0.001; NS > 0.05, unpaired Student’s t-test.

These findings offered valuable input for exploring the possible integration of pCharme functions into pathways controlling the maturation of cardiomyocytes. In this direction, we analysed *in silico* the lncRNA promoter to search for transcription factors (TFs) acting as upstream regulators of pCharme expression in cardiomyocytes. In accordance with our previous findings (Ballarino et al., 2015), Jaspar database (https://jaspar.genereg.net; Castro-Mondragon et al., 2022) identified several MYOD1 *consensus* sites in the 1kb region upstream of pCharme transcriptional start site (TSS, **Figure 1F**), although it is known that this myogenic regulator is not expressed in the heart (Olson, 1993; Buckingham et al., 2017). To note, a very low number of *consensus* sites was found for ONECUT2, a TF involved in neurogenesis (Aydin et al., 2019). Searching for cardiac regulators, we identified canonical motifs (**Figure 1-figure supplement 1F**) and enriched sites (**Figure 1F**) for the T-box transcription factor-5 (TBX5). In addition to being a key regulator of heart development, known to activate genes associated to CM maturation in early development and morphogenesis at later stages (Nadadur et al., 2016; Steimle et al., 2017), TBX5 was recently found to control the expression of several lncRNAs (Yang et al., 2017), a subset of them intriguingly enriched, as for pCharme, in the chromatin fraction of cardiomyocytes (Hall et al., 2021). By further analysing the scRNA-seq dataset (Jackson-Weaver et al., 2020), we observed a positive and highly significant correlation between *Tbx5* and pCharme (**Figure 1G**, left) in CM, with some differences occurring across the CM cluster subtypes (see for instance CM-IVS/CM-OFT, **Figure 1G**, right). To note, pCharme positively correlates with markers of cardiomyocyte identity, Myosin light chain 2 (*Myl2*) and 4 (*Myl4*), while significant anticorrelation was found for the Kruppel like factor 2 (*Klf2*) and the Twist family BHLH transcription factor 1 (*Twist1*) transcripts, that respectively mark endothelial and neural crest cells (Jackson-Weaver et al., 2020; Li et al., 2016).

To provide functional support to a possible role of TBX5 for pCharme transcription in CM, we examined available Chromatin Immunoprecipitation (ChIP)-sequencing (**Figure 1H**) and RNA-seq (**Figure 1I**) datasets from murine cardiomyocytes at progenitor (P-CM) and differentiated (D-CM) stages (Luna-Zurita et al., 2016). In both conditions, ChIP-seq analyses revealed the specific occupancy of TBX5 to highly conserved spots within the pCharme promoter, which increases with differentiation. However, while in differentiated cardiomyocytes pCharme expression was insensitive to TBX5 ablation, the lncRNA abundance was consistently decreased in TBX5 knockout progenitor cells (**Figure 1I**). As progenitor CM represent a cell resource for recapitulating *ex-vivo* the early establishment of the cardiac lineage and for mimicking cardiomyocyte development (Kattman et al., 2011), these results are consistent with the role of TBX5 for Charme transcription at fetal stages. The presence of *consensus* sites for other TFs (**Figure 1-figure supplement 1F)** and the known attitude of TBX5 to interact, physically and functionally, with several cardiac regulators (Akerberg et al., 2019), also predict the possible contribution of other regulators to pCharme transcription which may act, in cooperation or not, with TBX5 during CM maturation.

### Genome-wide profiling of cardiac Charme^WT^ and Charme^KO^ transcriptomes

Over the years, we have accumulated strong evidence on the role played by pCharme in skeletal myogenesis (Ballarino et al., 2018; Desideri et al., 2020), however important questions are still pending on the role of pCharme in cardiomyogenesis. As a first step towards the identification of the molecular signature underlying Charme-dependent cardiac anomalies, we performed a differential gene expression analysis on transcriptome profiles from Charme^WT^ and Charme^KO^ neonatal (PN2) hearts (**Figure 2A**; **Figure 2-figure supplement 1A**). At this stage, the heart continues to change undergoing maturation, a process in which its morphology and cell composition keep evolving (Padula et al., 2021; Tian et al., 2017).

**Figure 2.**
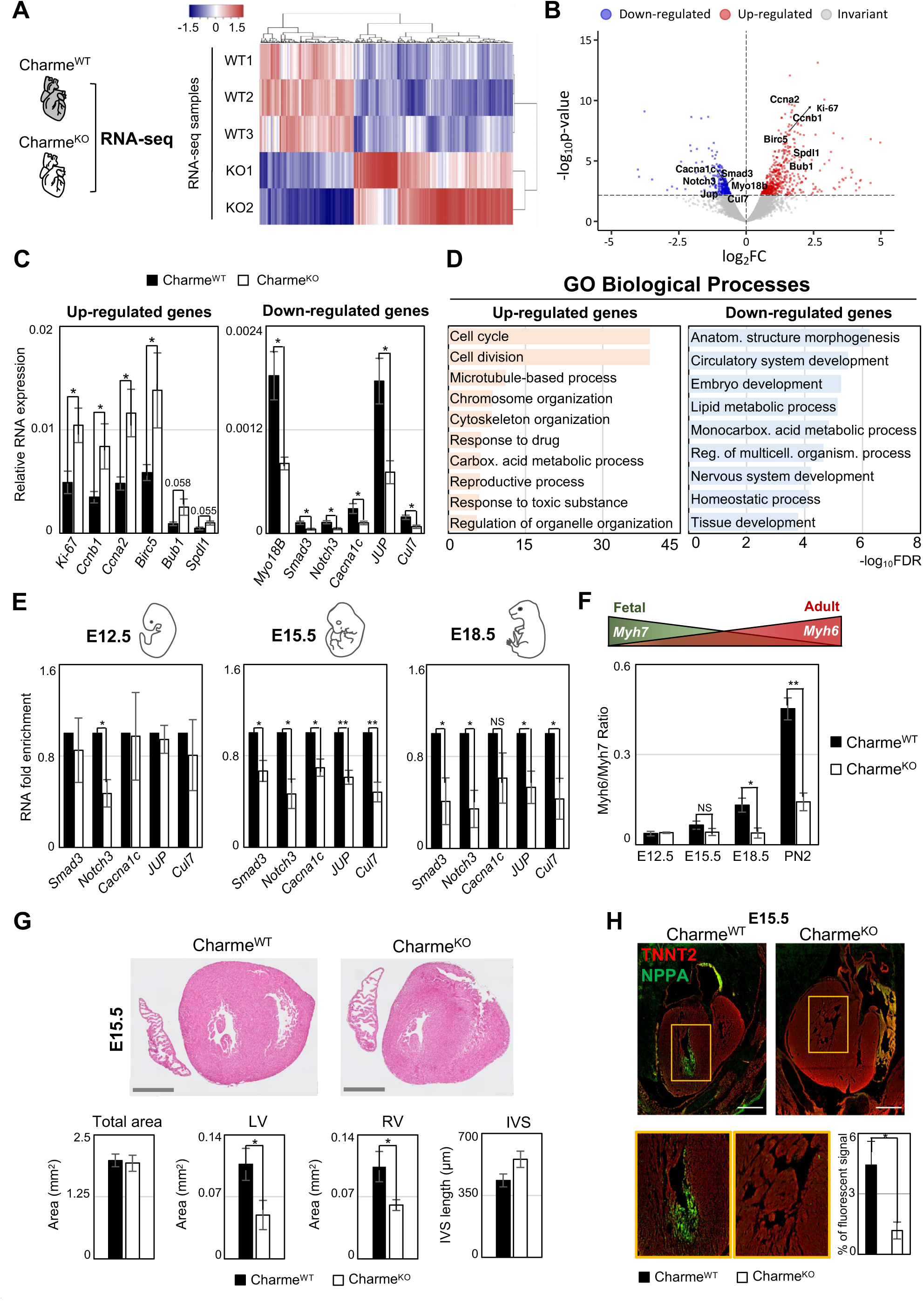
Genome-wide profiling of cardiac Charme^WT^ and Charme^KO^ transcriptomes. **A)** Heatmap visualization from RNA-seq analysis of Charme^WT^ and Charme^KO^ neonatal (PN2) hearts. Plot was produced by heatmap3 (https://cran.r-project.org/web/packages/heatmap3/vignettes/vignette.pdf). Expression values were calculated as FPKM, were log_2_-transformed and mean-centered. FPKM: Fragments Per Kilobase of transcript per Million mapped reads. **B)** Volcano plots showing differential gene expression from transcriptome analysis of Charme^WT^ *vs* Charme^KO^ PN2 hearts. Differentially expressed genes (DEGs) validated through RT-qPCR (Figure 2C) are in evidence. FC: Fold Change **C)** RT-qPCR quantification of Up-regulated (left panel) and Down-regulated (right panel) DEGs in Charme^WT^ *vs* Charme^KO^ neonatal hearts. Data were normalized to *Gapdh* mRNA and represent means ± SEM of WT (n=5) *vs* KO (n=4) pools **D)** Gene Ontology (GO) enrichment analysis performed by WEBGESTALT (http://www.webgestalt.org) on Up-regulated (left panel) and Down-regulated (right panel) DEGs in Charme^WT^ *vs* Charme^KO^ pools of neonatal hearts. Bars indicate the top categories of Biological processes in decreasing order of – log_10_FDR. All the represented categories show a False Discovery Rate (FDR) value <0.05. **E)** RT-qPCR quantification of pCharme targets in Charme^WT^ and Charme^KO^ extracts from E12.5, E15.5 and E18.5 hearts. DEGs belonging to the GO category “anatomical structure morphogenesis” were considered for the analysis. Data were normalized to *Gapdh* mRNA and represent means ± SEM of WT and KO (n=3) pools. **F)** RT-qPCR quantification of the Myh6/Myh7 ratio in Charme^WT^ and Charme^KO^ extracts from E12.5, E15.5 and E18.5 and neonatal hearts. Data were normalized to *Gapdh* mRNA and represent means ± SEM of WT and KO (n=3) pools. Schematic representation of the physiological *Myh6*/*Myh7* expression trend is shown. **G)** Upper panel: Haematoxylin-eosin staining from Charme^WT^ and Charme^KO^ E15.5 cardiac transverse sections. Scale bars: 500 μm. Lower panel: Quantification of the total area, the left and right ventricle cavities and the thickness of the interventricular septum (IVS) in Charme^WT^ and Charme^KO^ E15.5 hearts. For each genotype, data represent the mean ± SEM (n=3). **H)** Representative images of Nppa (green) and TnnT2 (red) immunostaining in Charme^WT^ and Charme^KO^ (E15.5) cardiac sections. Regions of interest (ROI, orange squares) were digitally enlarged on the lower panels. Scale bar: 500 µm. Quantification of the area covered by the Nppa fluorescent signal is shown aside. Data represent the mean (%) ± SEM of WT (n=4) and KO (n=3). Data information: *p < 0.05; **p < 0.01, NS > 0.05, unpaired Student’s t-test

RNA-seq analysis of whole hearts led to the identification of 913 differentially expressed genes (DEGs) (FDR<0.1, WT *vs* KO, **Supplementary File 2**), 573 of which were Up-regulated and 340 were Down-regulated in Charme^KO^ hearts (**Figure 2B**; **Supplementary File 2**). These results were confirmed by RT-qPCR analyses performed on gene subsets from independent biological replicates (**Figure 2C**; **Figure 2-figure supplement 1B**). Similar to pCharme activity in skeletal myocytes (Ballarino et al., 2018), we did not find DEGs within the neighbouring chromatin environment (370 Kb around Charme) (**Figure 2-figure supplement 1C**).

A Gene Ontology (GO) term enrichment study was then applied separately to the Up- and Down-regulated DEGs. These analyses revealed that the Up-regulated DEGs were primarily enriched (FDR-values < 1,0E-10) in cell cycle and cell division categories (**Figure 2D**, left panel), which parallels with a slight increase in the number of KI-67^+^ mitotic Charme^KO^ nuclei, as quantified from neonatal cardiac sections (**Figure 2-figure supplement 1D**). The same study applied to the Down-regulated DEGs revealed their enrichment to more functional and morphogenetic GO categories and primarily referred to anatomical structure morphogenesis (FDR 5.77E-07) and circulatory system development (FDR 3.1338E-06) (**Figure 2D**, right panel). Interestingly, these top-ranked categories include TFs involved in pivotal steps of embryo development, such as *Smad3* (Dunn et al., 2004) and *Nocth3* (MacGrogan et al., 2018), and functional components of cardiomyocytes, such as the voltage-dependent calcium channel subunit *Cacna1c* (Wang et al., 2018), and Myosin-18B (*Myo18B*), known to regulate cardiac sarcomere organisation (Latham et al., 2020). By expanding the analysis of these genes from neonatal to embryonal timepoints (E12.5, E15.5 and E18.5), we found that their expression significantly decreases upon pCharme ablation from the fetal E15.5 stage onward, as compared to WT (**Figure 2E**). Similarly, the *Myh6*/*Myh7* ratio, which measures CM maturation (England and Loughna, 2013; Scheuermann and Boyer, 2013) and that increases over the wild-type timepoints, displays a gradual decrease in the Charme^KO^ hearts from E15.5 onward (**Figure 2F**). To note, the expression of cell-cycle and proliferation genes was not affected by pCharme ablation over the same developmental window (**Figure 2-figure supplement 1E**). Hence, their upregulation in neonatal KO hearts could be due to the impairment of cardiac maturation, which leads to a shift to a more fetal and proliferative state of cardiomyocytes upon birth.

Overall, our findings suggest that during cardiac development, pCharme regulation is achieved through the regulation of morphogenetic pathways whose dysregulation causes a pathological remodelling of the heart. Accordingly, histological analyses performed by haematoxylin and eosin staining of Charme^WT^ and Charme^KO^ cardiac cryo-sections (E15.5) showed a pronounced alteration of the myocardium, with a clear decrease of the area of the ventricular cavities (**Figure 2G**). The histological investigation was then deepened to the trabeculated myocardium, the tissue directly surrounding ventricle cavities. To this purpose, we performed immunofluorescence staining and western blot analysis for the natriuretic peptide A (NPPA) factor, a known marker of the embryonal trabecular cardiomyocytes (Choquet et al., 2019; Horsthuis et al., 2008). In parallel, heart sections were analysed for cardiac vasculature, since i) the process of trabeculae formation is known to be temporally coupled with the formation of blood vessels during development (Samsa et al., 2013) and, importantly, ii) the “circulatory system development” category was among the main GO enriched for pCharme down-regulated targets. We found that, with respect to WT, mutant hearts display a significant reduction of NPPA^+^ trabeculae (**Figure 2H; Figure 2-figure supplement 1F**), which parallels the decreased expression of the *Irx3* and *Sema3a* trabecular markers (Choquet et al., 2019) (**Figure 2-figure supplement 1G**) as well as the diminished density of the capillary endothelium, as imaged by lectin staining (**Figure 2-figure supplement 1H**). Therefore, the fetal expression of pCharme is necessary for the achievement of morphogenetic programs important for CM maturation, myocardial geometry and vascular network formation. Overall, these ensure the preservation of the cardiac function and structure in adulthood, as a progressive deterioration of the systolic function, which becomes significant at 9 months of age (**Figure 2-figure supplement 1I**; **Supplementary File 3**), was observed in Charme^KO^ mice. To note, a similar alteration in heart efficiency was observed in the murine model with a specific mutation in the pCharme intron-1 (Charme^ΔInt^) (Desideri et al., 2020), further confirming the distinct activity of this isoform in cardiac processes.

### pCharme nucleates the formation of RNA-rich condensates by interacting with MATR3 in fetal cardiomyocytes

In differentiated myocytes, we previously demonstrated that CU-rich binding motifs inside pCharme coordinate the co-transcriptional recruitment and the subcellular localization of Matrin 3 (MATR3) (Desideri et al., 2020), a nuclear matrix RNA/DNA binding protein involved in multiple RNA biosynthetic processes (Coelho et al., 2016; Banerjee et al., 2017) and recently shown to play a role in chromatin repositioning during development (Cha et al., 2021). Interestingly, MATR3 was found highly expressed in cardiomyocytes from newborn mice and heterozygous mutations in MATR3 resulted in congenital heart defects (Quintero-Rivera et al., 2015). This raises the intriguing possibility that pCharme may play a role in cardiomyocyte maturation through MATR3. To examine this hypothesis, we first assessed the subcellular localization of pCharme in fetal (E15.5) hearts by biochemical fractionation. RT-qPCR analyses revealed that, in line with what was previously observed in skeletal muscle, in cardiomyocytes pCharme mainly localises in the nucleus, while the fully spliced mCharme is enriched in the cytoplasm (**Figure 3-figure supplement 1A**). We next applied high-resolution RNA-fluorescence *in situ* hybridization (RNA-FISH) to visualise pCharme alone (**Figure 3A**; **Figure 3-figure supplement 1B**) or relatively to MATR3 through a combined immunofluorescence IF/RNA-FISH approach (**Figure 3B**). In agreement with the subcellular fractionation, the imaging experiments confirmed the nuclear localization of pCharme, which exhibits its typical punctuate pattern. Further analysis of the three-dimensional distribution of pCharme and MATR3 revealed a clear colocalization of their signals and the formation of nuclear condensates, as quantified by 3D Pearson’s correlation coefficient applied on the overlapping signals (**Figure 3-figure supplement 1C**). Based on these results, we then tested if the presence of pCharme could influence the nuclear localization of MATR3. To this end, we performed MATR3 IF assays in wild-type and Charme^KO^ fetal (E15.5) muscle biopsies and in spinal cord nuclei (**Figure 3C**; **Figure 3-figure supplement 1D-F**). A striking heterogeneous distribution of MATR3 positive signals was observed within the nucleus of wild-type skeletal (**Figure 3-figure supplement 1D**) and cardiac (**Figure 3C**, upper panel) muscles, both expressing the lncRNA. Coherently with their pCharme-dependent formation, these condensates appeared more diffuse in tissues where the lncRNA is not expressed, as Charme^KO^ muscles and WT spinal cords (**Figure 3C**, middle and lower panels; **Figure 3-figure supplement 1D**). More accurate quantification of the MATR3 fluorescence distribution in the nucleus of Charme^KO^ cardiomyocytes (Coefficient of variation (CV), **Figure 3-figure supplement 1F**), revealed a MATR3 IF pattern which was less discrete and more homogeneous in respect to WT.

**Figure 3.**
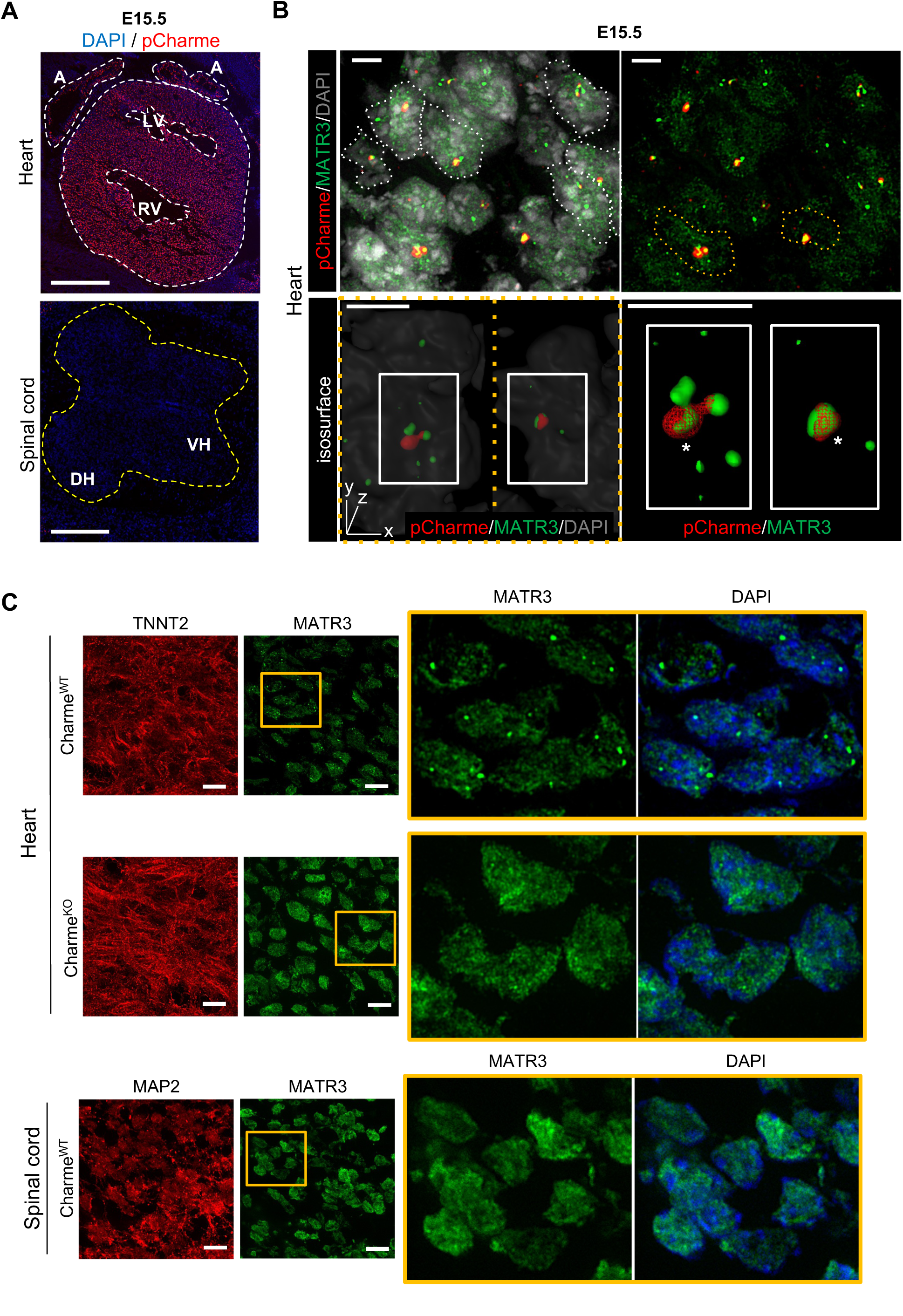
In fetal cardiomyocytes pCharme nucleates the nuclear localization of MATR3. **A)** RNA-FISH for pCharme (red) and DAPI staining (blue) in Charme^WT^ cardiac and spinal cord from E15.5 tissue sections. Whole heart (white dashed lines), spinal cord (yellow dashed line). A: Atria; LV and RV: Left and Right ventricle; DH and VH: Dorsal and Ventral Horn. Scale bars, 500 μm. **B)** Upper panel: RNA-FISH for pCharme (red) combined with immunofluorescence for MATR3 (green) and DAPI staining (gray) in Charme^WT^ from E15.5 cardiac sections. Dashed lines show the edge of nuclei. Lower panel: selected nuclei (yellow dashed lines in the upper panel) were enlarged and processed for isosurface reconstruction (left panel) and digital magnification (right panel). Overlapped signals are shown (asterisks). Scale bars, 5 μm. **C)** Upper panel: Representative images for MATR3 (green), TnnT2 (red) and DAPI (blue) stainings on Charme^WT^ and Charme^KO^ E15.5 cardiac sections. Lower panel: Representative images for for MATR3 (green), Map2 (red) and DAPI (blue) stainings on Charme^WT^ and Charme^KO^ E15.5 spinal cord sections. ROI (orange squares) were digitally enlarged on the right panels. Each image is representative of three individual biological replicates. Scale bars, 10 μm.

The known ability of MATR3 to interact with both DNA and RNA and the high retention of pCharme on the chromatin may predict the presence of chromatin and/or specific transcripts within these MATR3-enriched condensates. In skeletal muscle cells, we have previously observed on a genome-wide scale, a global reduction of MATR3 chromatin binding in the absence of pCharme (Desideri et al., 2020). Nevertheless, the broad distribution of the protein over the genome made the identification of specific targets through MATR3-ChIP challenging. With the purpose to functionally characterise MATR3 interactome, here we applied a more straightforward MATR3 cross-linking immunoprecipitation (CLIP)-seq approach to fetal Charme^WT^ and Charme^KO^ hearts for the identification of MATR3 bound RNA targets (**Figure 4A**). Western blot analyses with antibodies against MATR3 allowed to test the efficiency of protein recovery after the immunoprecipitation, in both the WT and KO conditions (**Figure 4-figure supplement 1A**). Subsequent analysis of the RNA, led to the identification of 951 cardiac-expressed transcripts significantly bound by MATR3 in the WT heart (log2 Fold enrichment >2 and FDR value <0.05, **Figure 4-figure supplement 1B** and **Supplementary File 4**). Four candidates (*Cacna1c*, *Myo18B*, *Tbx20*, *Gata4*) were selected based on their FDR values (**Supplementary File 4**) for further validation by RT-qPCR, together with *Gapdh* as a negative control (**Figure 4-figure supplement 1C**).

**Figure 4.**
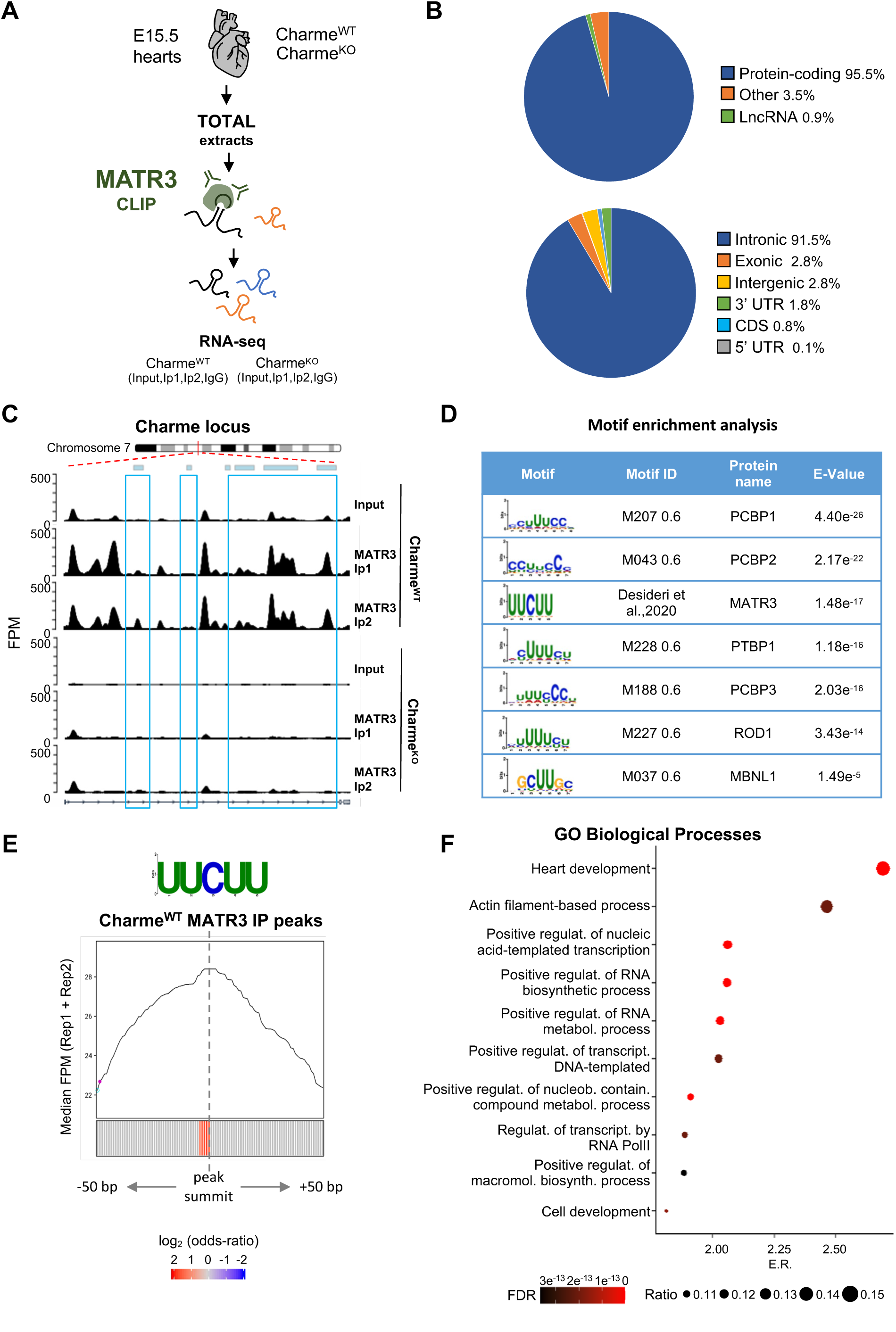
MATR3/pCharme nuclear condensates contain key regulators of heart development. **A)** Schematic representation of MATR3 CLIP-seq workflow from fetal (E15.5) Charme^WT^ and Charme^KO^ hearts. See **Materials and Methods** for details. **B)** MATR3 CLIP-seq from fetal hearts. Upper panel: a pie-plot projection representing transcript biotypes of 951 identified MATR3 interacting RNAs. Peaks overlapping multiple transcripts were assigned with the following priority: protein coding, lncRNA and others. Lower panel: a pie-plot projection representing the location of MATR3 enriched peaks (log2 Fold enrichment > 2 and FDR < 0.05). Peaks overlapping multiple regions were assigned with the following priority: CDS, 3’UTR, 5’UTR, exons, introns and intergenic. Percentages relative to each group are shown. **C)** MATR3 CLIP-seq (Input, Ip1 and Ip2) normalized read coverage tracks (FPM) across pCharme from fetal hearts. Significant MATR3 peaks, displaying log_2_ Fold enrichment > 2 in both Ip1 and Ip2 samples compared to Input, are demarcated by light-blue boxes. Normalized read coverage tracks (FPM) from MATR3 CLIP-seq in Charme^KO^ fetal hearts on Charme locus are also shown. Plot obtained using Gviz R package. **D)** Motif enrichment analysis perfomed on MATR3 CLIP-seq peaks (Charme^WT^) with AME software using 93 RNA binding motifs from CISBP-RNA database. 7 consensus motifs resulted significantly over-represented (Evalue < 0.05) among the MATR3 peaks, as compared to control regions. See **Material and Methods** for details. **E)** Positional enrichment analysis of MATR3 motif in MATR3 CLIP-seq peaks (Charme^WT^). For each analyzed position close to peak summit, line plot displays the median CLIP-seq signal (FPM, IP1 + IP2), while heatmap displays the log2 odds-ratio of UUCUU motif enrichment. Significant enrichments (p-value < 0.05) are shown in red. See **Material and Methods** for details. **F)** GO enriched categories obtained with WebGestalt (http://www.webgestalt.org) on protein-coding genes overlapping Charme^WT^ MATR3 peaks. Dots indicate the top categories of biological processes (description in y-axis) in decreasing order of Enrichment Ratio (E.R.= overlapped genes/expected genes, x-axis). Dot size (Ratio) represents the ratio between overlapped gens and GO categories size while dot color (FDR) represents significance. All the represented categories show an FDR<0.05.

In line with the binding propensities exhibited by MATR3 in other cell systems (Uemura et al., 2017), most of its enrichment was located within the introns of protein-coding mRNA precursors (**Figure 4B**). Strikingly, even though the class of lncRNAs was poorly represented (**Figure 4B**, upper panel), we found pCharme at the top of the MATR3 interactors in Charme^WT^ hearts (**Supplementary File 4**). Quantification of the retrieved RNAs by RT-qPCR evidenced the specific enrichment of pCharme but not mCharme in the wild-type samples, which supports the distinctive binding of MATR3 to the nuclear isoform (**Figure 4-figure supplement 1D**). As expected, we observed a strong reduction of MATR3-CLIP signal on Charme RNA in Charme^KO^ samples (**Figure 4C**, **Supplementary File 4**). To gain further insight into the specificity of MATR3 binding, we proceeded with the analysis of the CLIP-identified peaks for the identification of RNA-binding proteins consensus sites, including MATR3. Motif enrichment analysis using 93 RNA-binding motifs catalogued by CISBP-RNA database (Ray D, et al., 2013) pinpointed the MATR3 consensus sequence (UUCUU, Desideri et al., 2020) among the most over-represented pyrimidine-enriched motifs (Evalue < 0.05) (**Figure 4D**). When analysed in respect to the intensity of the CLIP signals, the MATR3 motif was positioned at the peak summit (+/- 50 nt) (**Figure 4E**), close to the strongest enrichment of MATR3, further confirming a direct and highly specific binding of the protein to these sites. Upon this quality assessment, we finally performed a GO enrichment analysis on the 882 protein-coding transcripts directly bound by MATR3 in the WT heart and found “Heart development” as the most significantly enriched functional category (**Figure 4F**). Therefore, our results indicate the formation of MATR3-containing condensates inside the nucleus of cardiomyocytes which contain pCharme, as the most abundant lncRNA, and pre-mRNA encoding for important cardiac regulators.

### The pCharme/MATR3 interaction sustains developmental gene expression in fetal cardiomyocytes

On a transcriptome scale, we found that in the heart the expression of MATR3-bound RNAs was globally reduced by pCharme knockout (**Figure 5-figure supplement 1A**). Indeed, while MATR3 targets were under-represented among transcripts whose expression was unaffected (invariant) or increased (up-regulated) by pCharme ablation, 12% of the downregulated DEGs (41 out of 340) were bound by MATR3 (**Figure 5A**). To note, these common targets were significantly enriched across the top-three GO classes, previously defined for Charme^KO^ downregulated genes (**Figure 5B**). This evidence suggests that pCharme is needed to sustain the cardiac expression of MATR3 targets. To investigate the possible implication of the lncRNA for MATR3 binding to RNA, we proceeded with a differential binding (DB) analysis of MATR3 CLIP-seq datasets between Charme^WT^ and Charme^KO^ conditions (**Figure 5-figure supplement 1B,C**). Among the shared peaks, DB revealed that the main consequence of pCharme depletion was a significant decrease of MATR3 enrichment on the RNA (**Figure 5C**, Loss), while the increase of protein binding was observed on a lower fraction of peaks (**Figure 5C**, Gain). In line with the specificity for MATR3 binding, also in these sets of differentially bound regions, the MATR3 motif was positioned close to the summit of the peak (**Figure 5-figure supplement 1D**). By mapping these differentially bound regions to the MATR3 targets which are also responsive to pCharme ablation (DEGs), we found that while the gained peaks were equally distributed, the loss peaks were significantly enriched in a subset (20 out of 40, 50%) of down-regulated DEGs (**Figure 5D**, upper panel). Overall, these results suggest that cooperation between pCharme and MATR3 activities is needed for the expression of specific RNAs, which majority (13 out of 20) is known to play important roles in embryo development, anatomical structure morphogenesis and development of the circulatory system (**Figure 5D**, lower panel). Indeed, we found interesting candidates such as *Cacna1c*, *Notch3*, *Myo18B* and *Rbm20*, whose role in cardiac physiopathology was extensively studied (Goonasekera et al., 2012; Tao et al., 2017; Ajima et al., 2008; van den Hoogenhof et al., 2018).

**Figure 5.**
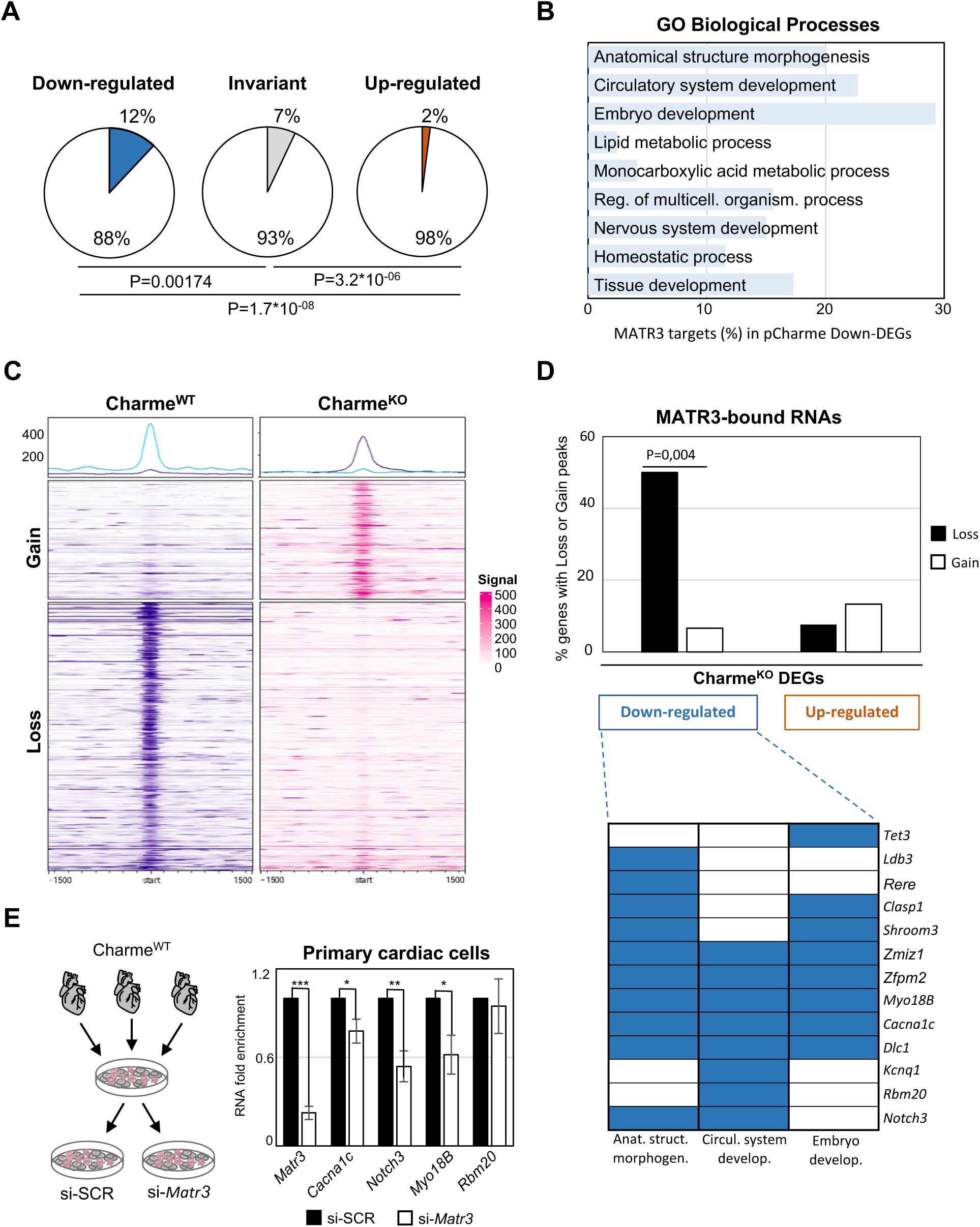
The pCharme/MATR3 interaction in cardiomyocytes sustains developmental genes expression. **A)** Pie charts showing the percentage of MATR3 targets in Charme^KO^ down-regulated, invariant or up-regulated DEGs. Significance of enrichment or depletion was assessed with two-sided Fisher’s Exact test, shown below. **B)** MATR3 targets (%) in the GO categories enriching Charme^KO^ down-regulated DEGs (Down-DEGs) (see also Figure 2A). **C)** Profile Heatmaps of differential MATR3 CLIP-seq peaks (Charme^WT^ *vs* Charme^KO^). Normalized mean read counts of both IP samples are shown only for significant (FDR < 0.05) «Gain» and «Loss» peaks. **D)** Upper panel: histogram showing the distribution (%) of «Gain» and «Loss» MATR3 peaks in pCharme DEGs. Significance of enrichment was assessed with two-sided Fisher’s Exact test. Lower panel: distribution of the subset (13 out of 20) of Down-DEGs with Loss peaks in the first three GO categories identified for down-regulated genes (see also Figure 2A). **E)** Left panel: Schematic representation of primary cells extraction from Charme^WT^ hearts. Once isolated, cells were plated and transfected with the specific siRNA (si-*Matr3*) or control siRNA (si-SCR). See Materials and Methods for details. Right panel: RT-qPCR quantification of *Matr3*, *Cacna1c*, *Notch3*, *Myo18B* and *Rbm20* RNA levels in primary cardiac cells treated with si-SCR or si-*Matr3*. Data were normalized to *Gapdh* mRNA and represent mean ± SEM of 4 independent experiments. Data information: *p < 0.05; **p < 0.01; ***p < 0.001, unpaired Student’s t test.

Finally, to univocally distinguish the contribution of pCharme and MATR3 on their expression, we tested the effect of MATR3 depletion on the abundance of these RNAs. We then purified cardiac primary cells from wild-type hearts for transfection with control (si-SCR) or MATR3 (si-*Matr3*) siRNAs (**Figure 5E**, left panel). RT-qPCR analysis revealed that 3 out of 4 tested genes exhibited a significant expression decrease upon MATR3 depletion (**Figure 5E**, right panel).

In sum, these data offer unprecedented knowledge on the RNA-binding propensities of MATR3 in the fetal heart and identify a subset of RNAs whose expression and MATR3 binding are influenced by pCharme. Our results demonstrate that in the developing heart RNA-rich MATR3-condensates form at the sites of pCharme transcription and control the expression of important regulators of embryo development, cardiac function, and morphogenesis. To the best of our knowledge, no previous research has given such an insight into the importance of specific lncRNA/RNA binding protein interactions occurring at the embryonal stages of mammalian heart development. As MATR3 is involved in multiple nuclear processes, further studies will be necessary to address how the early interaction of the protein with pCharme impacts on the expression of specific RNAs, either at transcriptional or post-transcriptional stages.

## DISCUSSION

In living organisms, the dynamic assembly and disassembly of distinct RNA-rich molecular condensates influences several aspects of gene expression and disease (Roden et al., 2021). The engagement of specific lncRNAs can enhance the biochemical versatility of these condensates because of the extraordinary tissue-specificity, structural flexibility and the propensity of this class of RNAs to gather macromolecules (Buonaiuto et al., 2021). Furthermore, the maintenance of specific lncRNAs on the chromatin combined with their scaffolding activity for RNA and proteins can cunningly seed high-local concentrations of molecules to specific loci (Bhat et al., 2021; Ribeiro et al., 2018). The occurrence of alternative RNA processing events eventually leads to the formation of diverse lncRNA isoforms, thus refining the biochemical properties and the binding affinities of these ncRNAs. This suggestive model perfectly fits with pCharme and provides mechanistic insights into the physiologic importance of this lncRNA in muscle. In fact, of the two different splicing isoforms produced by the Charme locus, only the unspliced and nuclear pCharme isoform was found to play an epigenetic, architectural function in skeletal myogenesis under physiological circumstances.

Here, we took advantage of our Charme^KO^ mouse model to extend the characterization of the lncRNA in embryogenesis. We show that the temporal and cell-type specific expression of pCharme is critical for the activation of pro-differentiation cardiac genes during development. Mechanistically, we provide evidence that this activity relies on the formation of pCharme-dependent RNA-rich condensates acting as chromatin hubs for MATR3, a nuclear matrix DNA/RNA-binding protein highly abundant in the fetal heart and source, upon mutation, of congenital defects (Quintero-Rivera et al., 2015). MATR3 ability to bind RNA can lead to the formation of dynamic shell-like nuclear condensates, whose composition depends on the integrity of the two RNA Recognition Motifs (RRM1 and 2) (Malik et al., 2018, Sprunger et al., 2022). Evidence supporting the physiological relevance of RNA-MATR3 interplay is emerging in both neural and muscle pathologies (Ramesh Net al., 2020; Senderek J, et al. 2009; Feit H, et al 1998.). In skeletal muscle, we previously showed that the presence of an intronic element bearing ∼100 MATR3 binding sites inside pCharme, is needed for MATR3 nuclear localization (Desideri et al., 2020). Here, we add an important dowel into the functional characterization of pCharme and give a developmental meaning to its interaction with MATR3 (**Figure 6**). We show that the absence of the lncRNA in cardiomyocytes alters the MATR3 nuclear distribution and its binding to specific RNAs, with a consequent mis-regulation of their developmental expression. Among them, we found a subset of shared pCharme/MATR3 targets, whose role in cardiac physiopathology was extensively studied, such as *Smad3*, *Notch3* and the myogenic components *Cacna1c* and *Myosin-18B* (Goonasekera et al., 2012; Tao et al., 2017; Ajima et al., 2008; van den Hoogenhof et al., 2018).

**Figure 6.**
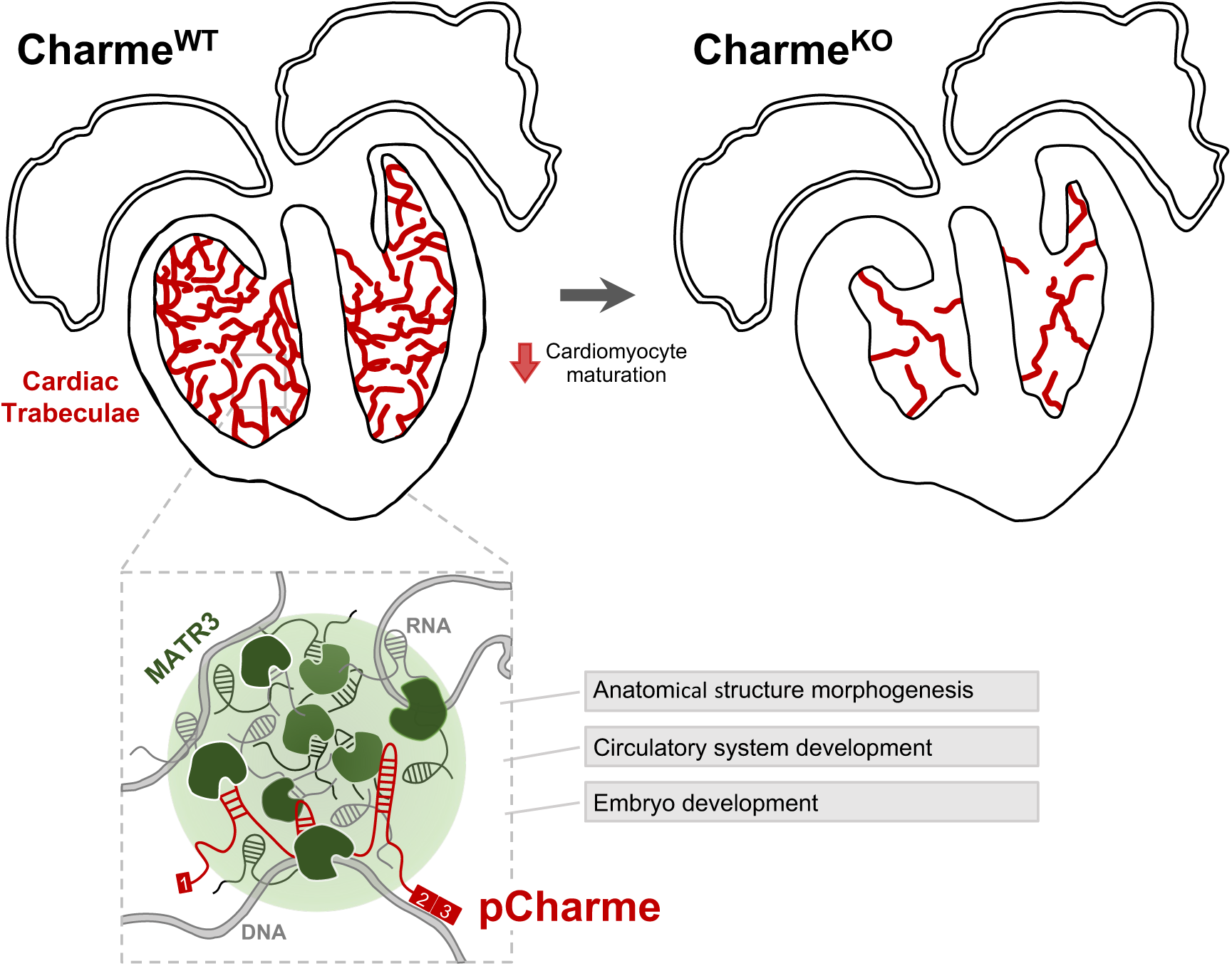
Proposed model for pCharme functions during heart development. At developmental stages (Charme^WT^), pCharme is required for the expression of genes involved in cardiomyocyte maturation. This activity is accompanied by the formation of nuclear condensates, through the interaction with the RNA-binding protein MATR3, which enrich transcripts involved in cardiac development. pCharme absence (Charme^KO^) leads to the reduction of cardiac trabeculae during development and to the remodelling of heart morphology.

It is interesting to note that, in the cardiac context, the reactivation of fetal-specific RNA-binding proteins, including MATR3, was recently found to drive transcriptome-wide switches through the regulation of early steps of RNA transcription and processing (D’Antonio et al., 2022). Along this view, the pCharme-dependent regulation of MATR3 binding in developing cardiomyocytes emphasises the need of cell-type specific lncRNAs for ruling the protein activities to specific RNAs, in a tissue-distinctive and spatio-temporal manner. As the impairment in the nuclear distribution of MATR3 can exert adverse effects in diverse tissues, our study may provide a more inclusive viewpoint on myopathies as well as other neuromuscular disorders where alteration in muscle development involves not only the myogenic lineage but also the interaction of the myogenic cells with the surrounding tissues.

These results also underlie the importance of studying cardiomyogenesis in animal models, which allow a better appreciation of the balance between the proliferation and maturation processes in cardiomyocytes and, importantly, the study of the cardiac functions in adulthood. In our case, adult Charme^KO^ mice develop a significant reduction of systolic function with initial signs of cardiac dilation, which denotes an early phase of cardiomyopathy. The results are supported by recently published data suggesting that the transition of cardiomyocytes toward an immature phenotype *in vivo* is associated with the development of dilated cardiomyopathy (Ikeda et al., 2019).

Recent cardiovascular studies have uncovered essential roles for lncRNAs in cardiac development and disease (Scheuermann and Boyer, 2013; Anderson et al., 2016; Ritter et al., 2019). However, a still unmet need is to disentangle non-canonical lncRNA-mediated mechanisms of action to gain insight into more successful diagnosis and classification of patient subpopulation but also to use them as possible diagnostic biomarkers or therapeutic targets (Buonaiuto et al., 2021). Future efforts will be devoted to clarifying the implication of the syntenic pCharme transcript in those human cardiomyopathies where pathological remodelling of the cardiac muscle occurs.

## MATERIALS AND METHODS

### Ethics Statement and animal procedures

C57BI/6J mice were used in this work. The Charme^KO^ animals were previously derived through the insertion of a PolyA signal in the Charme locus, as detailed in (Ballarino et al., 2018). All procedures involving laboratory animals were performed according to the institutional and national guidelines and legislations of Italy and according to the guidelines of Good Laboratory Practice (GLP). All experiments were approved by the Institutional Animal Use and Care Committee and carried out in accordance with the law (Protocol number 82945.56). All animals were kept in a temperature of 22°C ± 3°C with a humidity between 50% and 60%, in animal cages with at least 5 animals.

### Isolation, transfection, and subcellular fractionation of mouse primary heart cells

For primary heart cells isolation and transfection, 5 to 10 postnatal (PN3) hearts for each replicate (n=4) were pooled, harvested and kept at 37°C in culture medium (FBS 10%, 1X Non-Essential aminoacids, 1x PenStrep and DMEM high glucose). Hearts were mashed with pestles for 2 min and cell isolation performed according to manufacturer’s instructions (Neonatal Heart Dissociation Kit, Miltenyi Biotec). Cell suspension was centrifuged for 5 min at 600 x g and cells were resuspended in cell culture medium and plated in 22.1 mm plates. 1,5 million cells were transfected 48 hr later with 75 mM si-SCR or si-MATR3 in 3 µl/ml of Lipofectamine RNAiMAX (Thermo Fisher Scientific) and 100 µl/ml of Opti-MEM (Thermo Fisher Scientific), according to manufacturer’s specifications. Total RNA was collected 48 hr after transfection. See **Supplementary File 5** for siRNA sequences. Subcellular fractionation of primary embryonal (E15.5) cardiac cells was performed using the Paris Kit (Thermo Fisher Scientific, cat#AM1921), according to the manufacturer’s instructions.

### Whole-mount *in situ* hybridization

Embryos were fixed overnight in 4% paraformaldehyde (PFA) in phosphate-buffered saline (PBS) plus 0.1% Tween® 20 (PBT) at 4°C, dehydrated through a series of methanol/PBT solutions (25%, 50%, 75% and 100% methanol), and stored at -20°C until hybridization. Fixed embryos were rehydrated and rinsed twice in PBT. At this point, embryos were either digested with DNase and/or RNase or kept in PBT. All embryos were bleached in 6% hydrogen peroxide in PBT for 1h. Embryos were then rinsed 3 times in PBT for 5 min, digested with proteinase K (10 μg/ml in PBT) for 5 min at room temperature, washed once in 2 mg/ml glycine in PBT and twice in PBT for 5 min each, and post-fixed in 4% PFA/0.2% glutaraldehyde in PBT for 20 min at room temperature. Embryos were subsequently rinsed twice in PBT for 5 min and pre-hybridized at 70°C in hybridization solution (50% Formamide, 5x SSC, pH 5, 0.1% Tween 20, 50 μg/ml heparin, 50 μg/ml Torula RNA, 50 μg/ml salmon sperm DNA) for 2h. Embryos were then incubated overnight at 70°C in hybridization solution containing 500 ng/ml of denatured riboprobe. Riboprobes were generated by *in vitro* transcription in the presence of Digoxigenin-UTP (Roche Diagnostics). Antisense and sense Charme probes, used as specificity controls, were synthesised from linearized pBluescript-Charme_Ex2/3 plasmid. On the second day, embryos were washed twice in 50% formamide/4x SSC, pH 5/1% SDS and twice in 50% formamide/2x SSC, pH 5 for 30 min each at 55°C. Embryos were then rinsed three times for 5 min in MABT (100 mM maleic acid, 150 mM NaCl, pH 7.5, 0.1% Tween), blocked for 2 h at room temperature in 10% goat serum in MABT, and incubated overnight at 4°C in 1% goat serum in MABT with 1∶5000 alkaline phosphatase-coupled anti-Digoxigenin antibody (Roche Diagnostics). On the third day, embryos were washed in MABT twice for 5 min and 5 more times for 1 h each. Embryos were then rinsed twice in NTMT (100 mM NaCl, 100 mM Tris-HCl, pH 9.5, 50 mM MgCl2, 0.1% Tween) for 15 min each, followed by the staining reaction in BM Purple (Roche Diagnostics) in the dark for 30 min to 12 h. Stained embryos were fixed overnight in 4% PFA in PBT, stored in PBT and photographed under a stereomicroscope.

### Cryo-section *in situ* hybridization

Embryos were dissected in cold PBS (pH 7.4) and fixed in 4% w/v PFA for 24 h at 4°C. Following fixation, the embryos were cryoprotected either in 30% w/v sucrose in PBS (for PFA-fixed embryos) or in 30% w/v sucrose in 0.1 M Tris pH 7.5 (for Z7-fixed embryos), embedded in tissue freezing medium (Leica Microsystems), sectioned at 12 µm using a cryostat (Leica 1900UV) and transferred to superfrost plus (ROTH) slides. The sections were air-dried for at least 30 min and stored at −80°C until later use. For chromogenic detection, sections were post-fixed in 4% w/v PFA in PBS for 10 min or in Z7, washed three times in PBS (or twice in 0.1 M Tris–HCl pH:7, 0.05 M NaCl and once in PBS for Z7) and incubated in acetylating solution (1.3% v/v triethanolamine, 0.03 N HCl, 0.25% v/v acetic anhydrite) for 10 min. Sections were then washed in PBS, incubated in 1% v/v Triton-X-100 in PBS for 30 min and washed three times in PBS. Prehybridization was performed for 4-6 h in buffer H (50% v/v formamide, 5× SSC (0.75 M NaCl, 0.075 M sodium citrate), 5× Denhardt’s (0.1% bovine serum albumin, 0.1% and 0.1% Polyvinylpyrrolidone), 250 µg/ml yeast RNA and 500 µg/ml salmon sperm DNA). Hybridization was performed in a humidified chamber for 16 h at 65°C in H buffer with DIG-labeled probe added (400 µg/ml). The probes were generated by *in vitro* transcription in the presence of Digoxigenin-UTP (Roche Diagnostics). Following hybridization, sections were sequentially washed in 5× SSC (5 min, 65°C), 0.2× SSC (1 h, 65°C), 0.2× SSC (5 min, RT). Then they were incubated in AB buffer (0.1 M Tris pH 7.5, 0.15 M NaCl) for 5 min, and in blocking solution (10% v/v Fetal Calf Serum in AB) for 1–2 h at RT. Antibody incubation was performed for 16 h at 4°C in AB buffer supplemented with 1% v/v Fetal Calf Serum and anti-DIG antibody coupled to alkaline phosphatase (1∶5000 dilution; Roche). Sections were then washed thoroughly in AB and equilibrated in alkaline phosphatase buffer (AP - 0.1 M Tris–HCl pH: 9.5, 0.1 M NaCl, 0.05 M MgCl2) for 5 min. Alkaline phosphatase activity was detected in the dark in AP buffer supplemented with 45 mg/ml 4-nitrobluetetrazolium chloride (NBT, Roche) and 35 mg/ml 5-bromo-4-chloro-3-indolyl-phosphate (BCIP, Roche). The reaction was stopped with PBS and the sections were mounted in Glycergel (Dako). Sections were analysed and photographed under a stereomicroscope. Fluorescent detection was performed via Basescope™ assay (Advanced Cell Diagnostics, Bio-Techne) as previously described in D’Ambra et al., 2021, with little modifications according to the manufacturer’s instructions for tissue processing. Probes used to specifically detect pCharme RNA (ref. 1136321-C1) were custom produced by Advanced Cell Diagnostics and designed to specifically target the intronic sequence in order to detect the unspliced transcripts.

### Preparation of probe templates for *in-situ* hybridization experiments

pCharme exon 2 and exon 3 were PCR-amplified from cDNA extracted from myotubes using Charme_Up-BamHI and Charme_Down-EcoRI primers (**Supplementary File 5**). PCR products were cloned into pBluescript ks(-) upon BamHI and EcoRI (Thermo Fisher Scientific) enzymatic restriction.

### Histology

All hearts were fixed in 4% formaldehyde, embedded in OCT, and cut into 7 μm sections. After washing with PBS 3 times for 5 min, the sections were stained for 7 min with eosin (Merk, cat#109844). Subsequently, slides were washed 3 times with PBS and then incubated with haematoxylin (Merk, 105175) for 90 s.

### Immunohistochemistry

Fresh E15.5 and PN tissues were embedded in OCT and then frozen in isopentane pre-chilled in liquid nitrogen. Cryo-sections (10 μm of thickness) were fixed in PFA 4% at 4°C for 20 min prior staining with primary antibodies, as previously described. Antibodies and dilutions are reported in **Supplementary File 5**. DAPI, KI-67, NPPA, TNNT2 and G. simplicifolia lectin immunofluorescence signals (Figure 2H; Figure 2-figure supplement 1D,E,H) were acquired with Carl Zeiss Microscopy GmbH Imager A2 equipped with Axiocam503 color camera. MATR3, TNNT2 and MAP2 signals (Figure 3C; Figure 3-figure supplement 1D,E) were acquired as Z stacks (200 nm path) by inverted confocal Olympus IX73 microscope equipped with a Crestoptics X-LIGHT V3 spinning disk system and a Prime BSI Express Scientific CMOS camera. Images were acquired as 16bit 2048x2048 pixel file by using 100X NA 1.40 and 60X NA 1.35 oil (UPLANSApo) objectives and were collected with the MetaMorph software (Molecular Devices). The average number of KI-67 positive nuclei from n=4 Charme^WT^ and Charme^KO^ neonatal cardiac sections was determined by dividing the number of immunolabeled nuclei over the total number of nuclei in each microscope field. For each replicate, from 4 to 18 fields were analysed with ImageJ Software (Schneider C.A. et al., 2012). NPPA and G. simplicifolia lectins sub-tissutal staining (Figure 2H; Figure 2-figure supplement 1H) was analysed using ImageJ software (Schneider C.A. et al., 2012) and according to Fukuda et al., 2013, with little modifications. Briefly, we quantified the ratio between the area covered by the NPPA/lectins signal immunofluorescence signals and the total area of the left ventricle. To calculate the area, the left ventricle was selected with the line selection tool. A threshold was then applied to the selected region to measure the area covered by the signal. Plot in Fig. 2F represents the percentage of area covered by the NPPA/lectins signal divided by the area of the left ventricle.

Quantification of MATR3 puncta in Figure 3C was performed using ImageJ software (Schneider C.A. et al., 2012) according to (Higaki T. et al., 2020), with little modifications. Briefly, a “mask” from binarized images was created and used to measure the intensity of MATR3 fluorescence inside the nuclei and the standard deviation (SD). MATR3 distribution was evaluated by Coefficient Variation (CV) defined as CV= mean SD/mean MATR3 intensity. A higher CV value indicates a higher spatial variability of fluorescence distribution (punctate staining); conversely, a lower CV value corresponds to a higher uniformity of fluorescence distribution (diffuse staining). Statistical analysis was performed using t-test, and the differences between means were considered significant at P ≤ 0.05.

### RNA extraction and RT-qPCR analysis

Total RNA from cultured cells and tissues was isolated using TRI Reagent (Zymo Research), extracted with Direct-zol^TM^ RNA MiniPrep (Zymo Research), treated with DNase (Zymo Research), retrotranscribed using PrimeScript Reagent Kit (Takara) and amplified by RT-qPCR using PowerUp SYBR-Green MasterMix (Thermo Fisher Scientific), as described in Desideri et al., 2020. See Supplementary File 5 for oligos details.

### RNA-Seq Analysis

To reduce biological variability, Charme^WT^ and Charme^KO^ neonatal littermates (PN2) were sacrificed and hearts from the corresponding genotypes pooled together before RNA extraction (3 Charme^WT^ pools, 9 hearts each, 2 Charme^KO^ pools, 3 hearts each). Validation analyses were performed on 2 additional Charme^WT^ pools (6 hearts each) and 2 Charme^KO^ pools (4 hearts each). Principal component analysis (PCA) conducted on the RNA-seq data, revealed that the two groups were evidently distinguished for the first principal component (**Figure 2-figure supplement 1A**). Illumina Stranded mRNA Prep was used to prepare cDNA libraries for RNA-Seq that was performed on an Illumina Novaseq 6000 Sequencing system at IIT-Istituto Italiano di Tecnologia (Genova, Italy). RNA-seq experiment produced an average of 26 million 150 nucleotide long paired-end reads per sample. Dark cycles in sequencing from Novaseq 6000 machines can lead to a high quality stretches of Guaninines artifacts; in order to remove these artifacts, low quality bases and N stretches from reads were removed by Cutadapt software using “-u -U”,“--trim-n” and “—nextseq-trim=20” parameters (Martin et al., 2011). Illumina adapter remotion were performed using Trimmomatic software (Bolger et al., 2014). Reads whose length after trimming was <35 nt were discarded. Reads were aligned to GRCm38 assembly using STAR aligner software (Dobin et al., 2013). Gene loci fragment quantification was performed on Ensemble (release 87) gene annotation gtf using STAR – quantMode GeneCounts parameter. Read counts of “reverse” configuration files were combined into a count matrix file, which was given as input to edgeR (Robinson et al., 2010) R package for differential expression analysis, after removing genes with less than 10 counts in at least two samples. Samples were normalized using TMM. Model fitting and testing were performed using the glmFIT and glmLRT functions. Gene-level FPKM values were calculated using rpkm function from the edgeR package. FDR cutoff for selecting significant differentially expressed genes was set to 0.1. Genes with less than 1 average FPKM in both conditions were filtered out. Heatmap of differentially expressed genes was generated using heatmap3 R package (Zhao et al., 2014) from log2 transformed FPKM values. Volcano plot were generated using “Enhanced Volcano” R package (bioconductor.org/packages/release/bioc/vignettes/EnhancedVolcano/inst/doc/EnhancedVolcano). Gene Ontology analyses were performed on up-regulated and down-regulated protein coding genes using WebGestalt R package (Liao Y et al., 2019) applying Weighted Set Cover dimensionality reduction.

### Cross-linking immunoprecipitation (CLIP) assay

A total of 60 Charme^WT^ and 56 Charme^KO^ E15.5 embryonal hearts were collected, divided in two distinct biological replicates and pestled for 2 min in PBS, 1x PIC and 1x PMSF. For each replicate, the solution was filtered in a 70 μm strainer and the isolated cells were plated and UV-crosslinked (4,000 μJ) in a Spectrolinker UV Crosslinker (Spectronics corporation). Upon harvesting, cells were centrifuged 5 min at 600 x g and pellets resuspended in NP40 lysis buffer (50 mM HEPES pH 7.5, 150 mM KCl, 2 mM EDTA, 1 mM NaF, 0.5% (v/v) NP40, 0.5 mM DTT, 1x PIC,), incubated on ice for 15 min and sonicated at low intensity six times with Bioruptor^®^ Plus sonication device to ensure nuclear membrane lysis. Lysate was diluted to a final concentration of 1 mg/ml. 30 µl of Dynabeads Protein G magnetic particles (Thermo Fisher Scientific) per ml of total lysate were washed twice with 1 mL of PBS-Tween (0.02%), resuspended with 5 μg of MATR3 (Supplementary File 5) or IgG antibodies (Immunoreagents Inc.) and incubated for 2 h at room temperature. Beads were then washed twice with 1 mL of PBS-T and incubated with total extract overnight at 4°C. Beads were washed three times with 1 mL of HighSalt NP40 wash buffer (50 mM HEPES-KOH, pH 7.5, 500 mM KCl, 0.05% (v/v) NP40, 0.5 mM DTT, 1x PIC) and resuspended in 100 µl of NP40 lysis buffer. For RNA sequencing, 75 μl of the sample were treated for 30 min at 50°C with 1.2 mg/ml Proteinase K (Roche) in Proteinase K Buffer (100 mM Tris-HCl, pH 7.5, 150mM NaCl, 12.5 mM EDTA, 2% (w/v) SDS). For Western Blot analysis, 25 μl of the sample were heated at 95°C for 5 min and resuspended in 4x Laemmli sample buffer (BioRad)/50 mM DTT before SDS-PAGE.

### MATR3 CLIP-seq analysis

Trio RNA-Seq (Tecan Genomics, Redwood City, CA) has been used for library preparation following the manufacturer’s instructions. The sequencing reactions were performed on an Illumina Novaseq 6000 Sequencing system at IGA Technology services. CLIP-sequencing reactions produced an average of 25 million 150 nucleotide long paired-end reads per sample Adaptor sequences and poor quality bases were removed from raw reads using a combination of Trimmomatic version 0.39 (Bolger et al., 2014) and Cutadapt version 3.2 (Martin et al., 2011) softwares. Reads whose length after trimming was <35 nt were discarded. Alignment to mouse GRCm38 genome and Ensembl 87 transcriptome was performed using STAR aligner version 2.7.7a (Dobin et al., 2013). Alignment files were further processed by collapsing PCR duplicates using the MarkDuplicates tool included in the Picard suite version 2.24.1 (http://broadinstitute.github.io/picard/) and discarding the multi-mapped reads using BamTools version 2.5.1 (Barnett et al., 2011). Properly paired reads were extracted using SAMtools version 1.7 (Li et al., 2009). GRCm38 genome was divided into 200 bp long non-overlapping bins using the BEDtools makewindows tool (Quinlan et al., 2010). Properly paired fragments falling in each bin were counted using the BEDtools intersect tool filtering out reads mapping to rRNAs, tRNAs or mitochondrial genome in order to create sample-specific count files. These files were given as input to Piranha version 1.2.1 (Uren et al., 2012) using *–x -s –u 0* parameters to call significant bins for MATR3 Ip and IgG samples. BEDtools intersect was used to assign each genomic bin to genes using Ensembl 87 annotation. For each gene the bin signal distribution in the input sample was calculated after normalization of fragment counts by the total number of mapping fragments. Ip significant bins presenting normalized signals lower than the upper-quartile value of the related gene distribution were filtered out. After this filter, significant bins belonging to Ip samples were merged using BEDTools merge tool. The number of fragments overlapping identified peaks and the number overlapping the same genomic region in the Input sample were counted and used to calculate fold enrichment (normalized by total mapping fragments counts in each data set), with enrichment P-value calculated by Yates’ Chi-Square test or Fisher Exact Test where the observed or expected fragments number was below 5. Benjamini-Hochberg FDR procedure was applied for multiple testing corrections. Peaks presenting log2 fold enrichment over Input >2 and FDR < 0.05 in both Ip samples were selected as enriched regions (**Supplementary File 3).** BEDtools intersect tool was used to annotate such regions based on their overlap with Ensembl 87 gene annotation and to filter out transcripts hosting regions enriched in the IgG sample. Furthermore, htseq-count software (Anders et al., 2015) with *-s no -m union -t gene* parameters was used to count reads from deduplicated BAM files. Peaks overlapping transcripts with Input CPM (counts per million) > Ip CPM in both Ip samples were filtered out. Gene Ontology analysis was performed on protein coding genes overlapping enriched regions using WebGestalt R package (Liao et al., 2019). Bigwig of normalized coverage (RPM, reads per million) files were produced using bamCoverage 3.5.1 from deepTools tool set (http://deeptools.readthedocs.io/en/develop) on BAM files of uniquely mapping and deduplicated reads using *--normalizeUsing CPM* parameter (Ramirez et al., 2014). Normalized coverage tracks were visualized with IGV software (https://software.broadinstitute.org/software/igv/) and Gviz R package (Hahne et al., 2016).

Local Motif Enrichment analysis was performed by using 10 nt-long sliding windows. For each relative peak position/control set, the proportion of windows containing or not MATR3 motif was calculated and tested for significant differences using two tails Fisher exact test. CLIP-seq IP signal (FPM) relative to each position was retrieved from bigwig files using pyBigWig (https://github.com/deeptools/pyBigWig). Motif Enrichment Analysis was performed using AME software from MEME suite (Bailey et al., 2015) on a 100 nt window centred on the peak summit. For each identified peak, a control set was by selecting 10 random regions from expressed (CPM>0) RNA precursors. Differential analysis of MATR3 binding between Charme^WT^ and Charme^KO^ conditions was assessed using DiffBind software (Ross-Innes et al., 2012).

### Protein analyses

Protein extracts were prepared and analysed by western blot as in Desideri et al., 2020. See **Supplementary File 5** for antibodies details.

### Single cell transcriptomics

scRNAseq analysis was performed on publicly available datasets of E12.5 mouse hearts (SRR10969391, Jackson-Weaver et al. 2020). FASTQ reads were aligned, filtered and counted through the Cell Ranger pipeline (v4.0) using standard parameters. GRCm38 genome version was used in alignment step and annotation refers to Ensembl Release87. The dataset was cleaned (nFeature_RNA > 200 and < 6000, percent.mt > 0 and < 5, nCount_RNA > 500 and < 40000) and cells were clustered using Seurat 4.0.5 (Stuart et al., 2019). Cluster uniformity was then checked using COTAN (Galfrè et al., 2020) by evaluating if less than 1% of genes were over the threshold of 1.5 of GDI. If a cluster resulted not uniform, with more than 1% of genes above 1.5, a new round of clustering was performed. After this iterative procedure, the few remaining cells not fitting any cluster, were discarded. A dataset of 4014 cells and 34 clusters was obtained. COTAN function *clustersDeltaExpression* (github devel branch) was used to obtain a correlation coefficient and a *p*-value for each gene and each cluster. From the correlation matrix, a cosine dissimilarity between clusters was estimated and used to plot a dendrogram (with the ward.D2 algorithm, **Figure 1-figure supplement 1D**). The tree was used to decide which clusters could be merged. Cell type for each final cluster was assigned based on a list of markers (Jackson-Weaver et al., 2020; Li et al., 2016) as follows. Cardiomyocytes (1834 cells): Myh6+, Nppa+, Atrial CM (333 cells): are also Myl1+, Myl4+, Ventricular CM (1289): are also Myl2+, Myl3+, interventricular septum CM (117 cells): are also TBX20+, Gja5+ (Franco et al., 2006), Venous Pole CM (95 cells): also Osr1+ (Meilhac et al., 2018), Outflow Tract CM (72 cells): also Isl1+, Sema3c+, Neural crest cells (there are two clusters expressing their markers: 1.NC with 48 cells and 2.NC with 68 cells): Msx1+, Twist1+, Sox9+, Epicardial cells (359 cells): Cebpb+, Krt18+, Fibroblasts like cells (278 cells): Tcf21+, Fn1+, Endothelial cells (168 cells): Klf2+, Pecam1+, Cdh5+, Smooth muscle cells (710 cells): Cnn1+, Acta2+, Tagln2+, Tagln+, Hemopoietic myeloid cells (79 cells): Fcer1g+, Hemopoietic red blood cells (397 cells): Hba-a1+. The final UMAP plot with the cell assignment is shown in **Figure 1-figure supplement 1E**. Heatmap in **Figure 1-figure supplement 1F** shows a coherent assignment between final clusters and cell type with additional marker genes. For correlation analyses (**Figure 1G**), relevant genes in each subpopulation of interest (CM-VP, CM-OFT, A-CM, CM-IVS, V-CM) were analysed separately by applying COTAN on a test dataset composed by the subpopulation in exam, together with a fixed contrasting cell group with the SM and EC cells (see **Figure 1E**).

### Echocardiography

The echocardiographer was blinded to the phenotypes. Mice were anaesthetised with 2.5% Avertin (Sigma T48402) to perform echocardiographic structural (measurement of left ventricular diameters and wall thickness) and functional (fractional shortening) analyses with a VEVO 3100 (Visualsonics) using a mx550d probe. We used avertin since it does not induce significant cardiodepressant effects, potentially affecting our echocardiographic experiments compared to ketamine combinations, such as ketamine+xylazine. The fractional shortening (FS) of the left ventricle was calculated as FS% = (left ventricular end-diastolic diameter (LVEDD)-left ventricular end-systolic diameter (LVESD)/LVEDD) x 100, representing the relative change of the left ventricular diameters during the cardiac cycle. The mean FS of the left ventricle was determined by the average of FS measurements of the left ventricular contraction over 3 beats.

### Data accessibility

The data presented in this study will be openly available in NCBI Gene Expression Omnibus (GEO) database (https://www.ncbi.nlm.nih.gov/geo/), reference number GSE200878 for RNA-seq data and GSE200877 for MATR3 CLIP-seq data.

### Statistical methods and rigour

Statistical tests, p-values, and n for each analysis are reported in the corresponding figure legend. For each experiment, at least three individual animals or pools of littermates were used (see each figure legend for details). No sex bias was introduced by randomly choosing among male and female mice. All analyses were performed by 1 or more blinded investigators.

## Competing interest

The authors declare no competing interests.

Supplementary File 1. Charme Tss usage data collected from Zenbu genome browser

Supplementary File 2. RNA-seq in Charme^WT^ and Charme^KO^ neonatal hearts

Supplementary File 3. Ecocardiography measurement for Charme^WT^ and Charme^KO^ animals.

Supplementary File 4. MATR3 CLIP-seq in Charme^WT^ and Charme^KO^ fetal hearts

Supplementary File 5. List and sequences of the oligonucleotides, siRNAs, antibodies and imaging probes used.

## Acknowledgments

The authors acknowledge Pietro Laneve, Francesca Pagano, Andrea Cipriano and Marco D’Onghia for helpful discussion, Alessandro Calicchio for cloning the probe templates for *in-situ* hybridization analyses and Marcella Marchioni for technical help. This work was supported by grants from Sapienza University (prot. RM11916B7A39DCE5 and RM12117A5DE7A45B), POR FESR Lazio 2020-T0002E0001 and MUR PNRR “National Center for Gene Therapy and Drugs based on RNA Technology” (Project no. CN3221842F1B2436 CN3_Spoke 3) to MB.

---

**Figure 1-Figure supplement 1.**
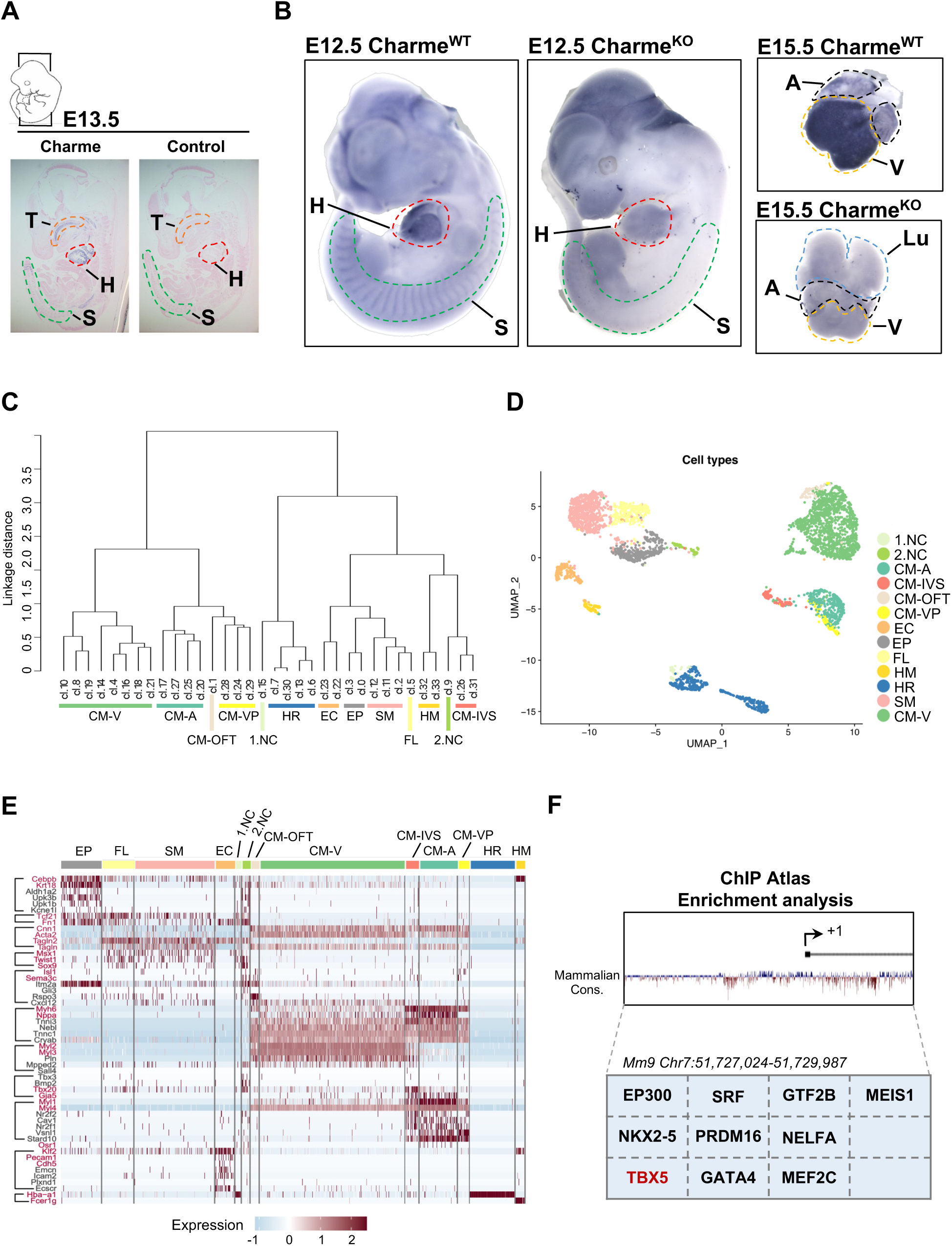
Related to Figure 1. **A)** *In-situ* hybridization (ISH) performed on E13.5 embryonal cryo-sections using digoxigenin-labelled RNA antisense (Charme, left panel) or sense (control, right panel) probes against Charme. T: Tongue (orange); H: Heart (red); S: Somites (green). **B)** Whole-mount *in-situ* hybridization (WISH) performed on Charme*^WT^* and Charme*^KO^* intact embryos (E12.5, left panels) and hearts (E15.5, right panels). Signal is specifically detected in heart (H, red line) and somites (S, green line). The specificity of the staining can be appreciated by the complete absence of signals in the Charme^KO^ samples. Heart (H, red line); Somites (S, green line); A: Atria (black line); V: Ventricles (yellow line). **C)** Dendrogram showing the relationships between homogeneous clusters. All the informative transcriptome was used to create a hierarchical clustering between homogeneous cell clusters. Colored lines mark which clusters were merged for the final clustering (Figure 1-Figure supplement 1D-E). CM: Cardiomyocytes, A-CM: Atrial-CM, V-CM: Ventricular-CM, ISV: Interventricular Septum, VP: Venous Pole, OFT: Outflow Tract, NC: Neural Crest cell, EP: Epicardial cells, FL: Fibroblasts like cells, EC: Endothelial Cells, SM: Smooth Muscle cells, HM: Hemopoietic Myeloid cells, HR: Hemopoietic Red blood cells. **D)** Seurat (Stuart et al., 2019) UMAP plot coloured by final cell assignments. See Materials and Methods for details. **E)** Heatmap was generated by Seurat DoHeatmap (Stuart et al., 2019) and represents, for each cell of the identified sub-populations, the log normalized expression of cell identity markers (listed on the left). Genes used for cell clustering are marked in red. Maximum expression value (red), minimum expression value (light blue). Correspondence between gene markers and cell types is indicated on the left. **F)** *In silico* analysis of cardiovascular TF binding sites on Charme promoter using the enrichment analysis tool on the ChIP Atlas database (https://chip-atlas.org/enrichment_analysis) by setting the threshold at 200.

**Figure 2-Figure supplement 1.**
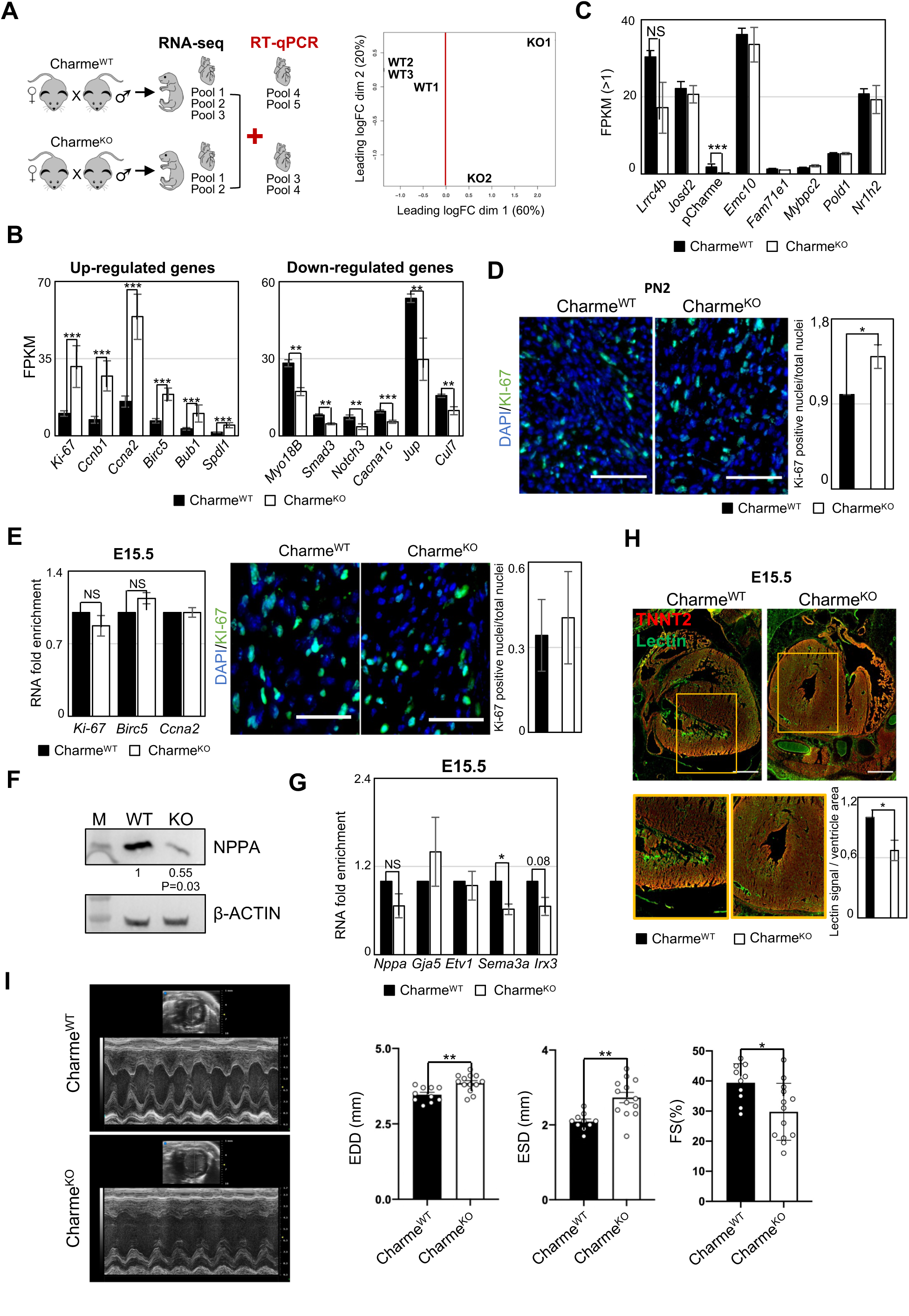
Related to Figure 2. **A)** Schematic overview of the workflow to identifying DEGs from Charme^WT^ and Charme^KO^ transcriptomes (left panel). Multi-dimensional scaling plot of leading fold-change (FC) between each pair of Charme^WT^ and Charme^KO^ RNA-seq samples (right panel). Plot was obtained by using the plotMDS function from edger package (https://www.bioconductor.org/packages/release/bioc/vignettes/edgeR/inst/doc/edgeRUsersGuide.pdf). **B)** Quantification by RNA-seq (FPKM) of Up-regulated (left panel) and Down-regulated (right panel) DEGs in Charme^KO^ *vs* Charme^WT^ neonatal hearts. **C)** Quantification by RNA-seq (FPKM) of pCharme and Charme-neighbouring genes expression in Charme*^WT^ vs* Charme^KO^ neonatal hearts. **D)** Left panel: Representative images for KI-67 (green) and DAPI (blue) stainings on Charme^WT^ and Charme^KO^ neonatal cardiac sections. Scale bars: 70 µm. Right panel: Quantification of KI-67 positive nuclei/total nuclei on Charme^WT^ and Charme^KO^ cardiac sections from neonatal mice. Data are expressed as mean ± SEM, n = 4. **E)** Left panel: RT-qPCR quantification of DEGs belonging to the “cell cycle” GO class from Charme^WT^ and Charme^KO^ E15.5 hearts. Data were normalized to *Gapdh* mRNA and represent means ± SEM of n=3 heart pools. Right panel: Representative images for KI-67 (green) and DAPI (blue) stainings on Charme^WT^ and Charme^KO^ E15.5 cardiac sections are shown. Quantification of KI-67 positive nuclei/total nuclei on Charme^WT^ and Charme^KO^ cardiac sections from E15.5 mice. Data are mean ± SEM.; n =3. Scale bars: 140 µm. **F)** Western blot analysis for NPPA in Charme^WT^ and Charme^KO^ E15.5 cardiac extracts. β-ACTIN was used as a loading control. Quantification of NPPA signal intensity relative to β-ACTIN is shown below. Data are mean ± SEM (n = 3). *p= 0.03. A representative image is shown. **G)** RT-qPCR quantification of trabeculae markers expression in Charme^WT^ *vs* Charme^KO^ E15.5 cardiac extracts. Data were normalized to *Gapdh* mRNA and represent means ± SEM of n=3 heart pools. **H)** Representative image of Lectin (green) and TnnT2 (red) immunostainings in Charme^WT^ and Charme^KO^ E15.5 cardiac sections. ROI (orange squares) were digitally enlarged on the lower panels. Scale bars: 500 µm. Quantification of the area covered by the Lectin fluorescence signal is shown aside. For each genotype, data represent the mean ± SEM of (at least) 6 images. n=3. **I)** Representative M-mode echocardiographic track of Charme^WT^ and Charme^KO^ 9-12 months aged mice. Quantification of heart morphology (EDD= end-diastolic diameter and ESD= end-systolic diameter) and function (FS: Fractional Shortening =(EDD-ESD)/EDD) were evaluated. Data represent the mean ± SEM (n = 10-13) Data information: *p < 0.05, **p < 0.01, ***p < 0.001, unpaired Student’s t test.

**Figure 3-Figure supplement 1.**
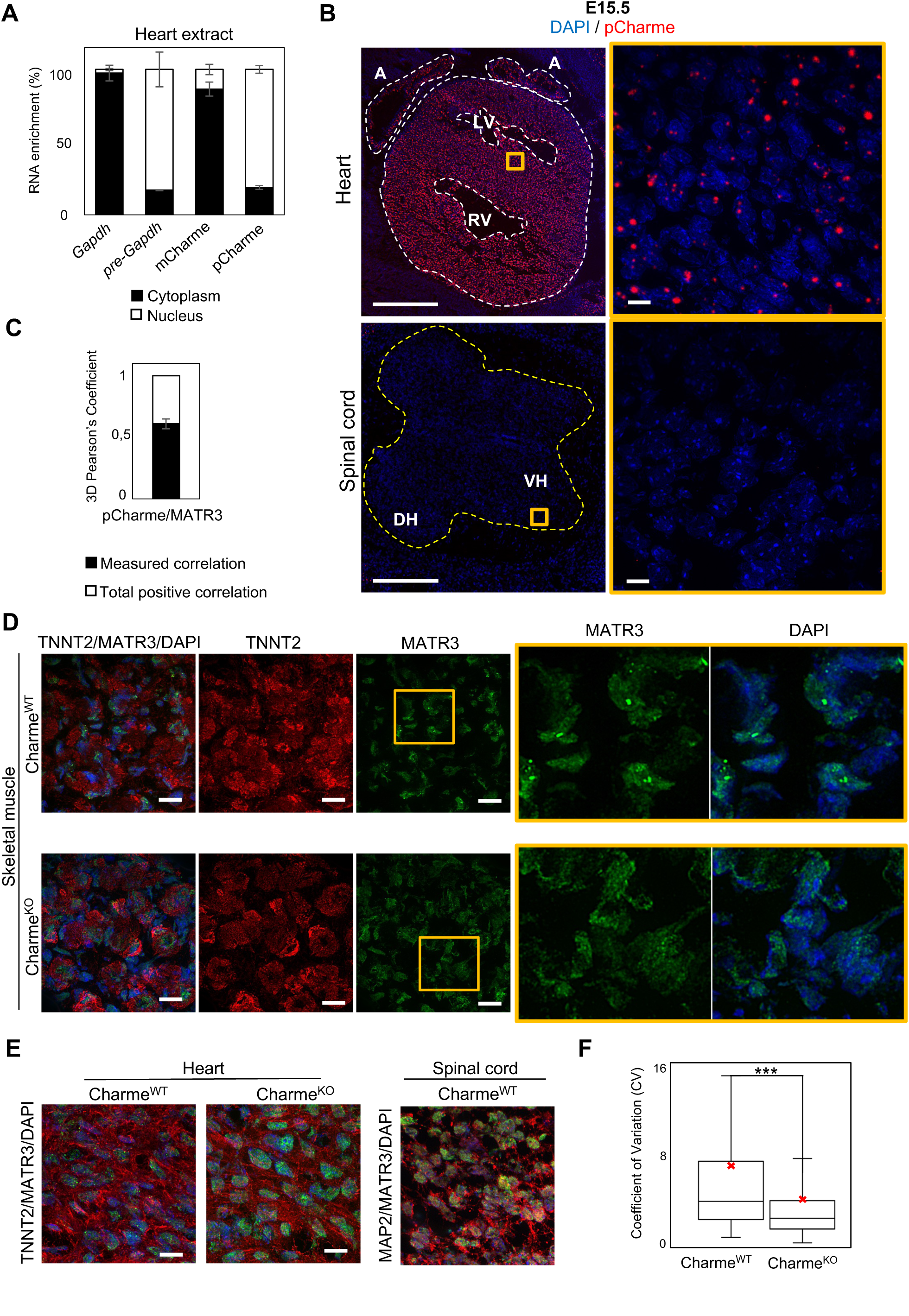
Related to Figure 3. **A)** Quantification of the subcellular distribution of pCharme and mCharme in cardiac tissues from neonatal mice. Histogram shows the quantification by RT-qPCR of the RNA abundance (%) in cytoplasmic *versus* nuclear compartments. *Gapdh* and *pre-Gapdh* RNAs were used, respectively, as cytoplasmic and nuclear controls. **B)** RNA-FISH for pCharme (red) and DAPI staining (blue) in Charme^WT^ hearts and spinal cord from E15.5 tissue sections (left panels) and their magnification (right panels). Whole heart (white dashed lines), spinal cord (yellow dashed line). A: Atria; LV and RV: Left and Right ventricle; DH and VH: Dorsal and Ventral Horn. Scale bars, 500 μm; 10 μm for magnifications. **C)** 3D Pearson’s correlation coefficient of pCharme/MATR3 overlapping signals. Histogram shows the mean ± SEM calculated over 237 colocalized nuclear signals from 3 independent experiments. **D)** Representative images for for MATR3 (green), TnnT2 (red) and DAPI (blue) stainings on Charme^WT^ and Charme^KO^ skeletal muscles from E15.5 tissue sections. ROI (orange squares) were digitally enlarged on the right panels. Each image is a representative of three individual samples. Scale bars, 10 μm. **E)** Left panel: Representative images for for MATR3 (green), TnnT2 (red) and DAPI (blue) stainings on Charme^WT^ and Charme^KO^ heart from E15.5 tissue sections. Right panel: Representative images for MATR3 (green), Map2 (red) and DAPI (blue) stainings on Charme^WT^ and Charme^KO^ spinal cord from E15.5 tissue sections. Each image is a representative of three individual samples. Scale bars, 10 μm. **F)** Quantification of MATR3 fluorescence intensity distribution (CV=Coefficient of variation) in Charme^WT^ (N=1346 nuclei) and Charme^KO^ (N=1404 nuclei). The red X indicate the mean value of CV distribution.

**Figure 4-Figure supplement 1.**
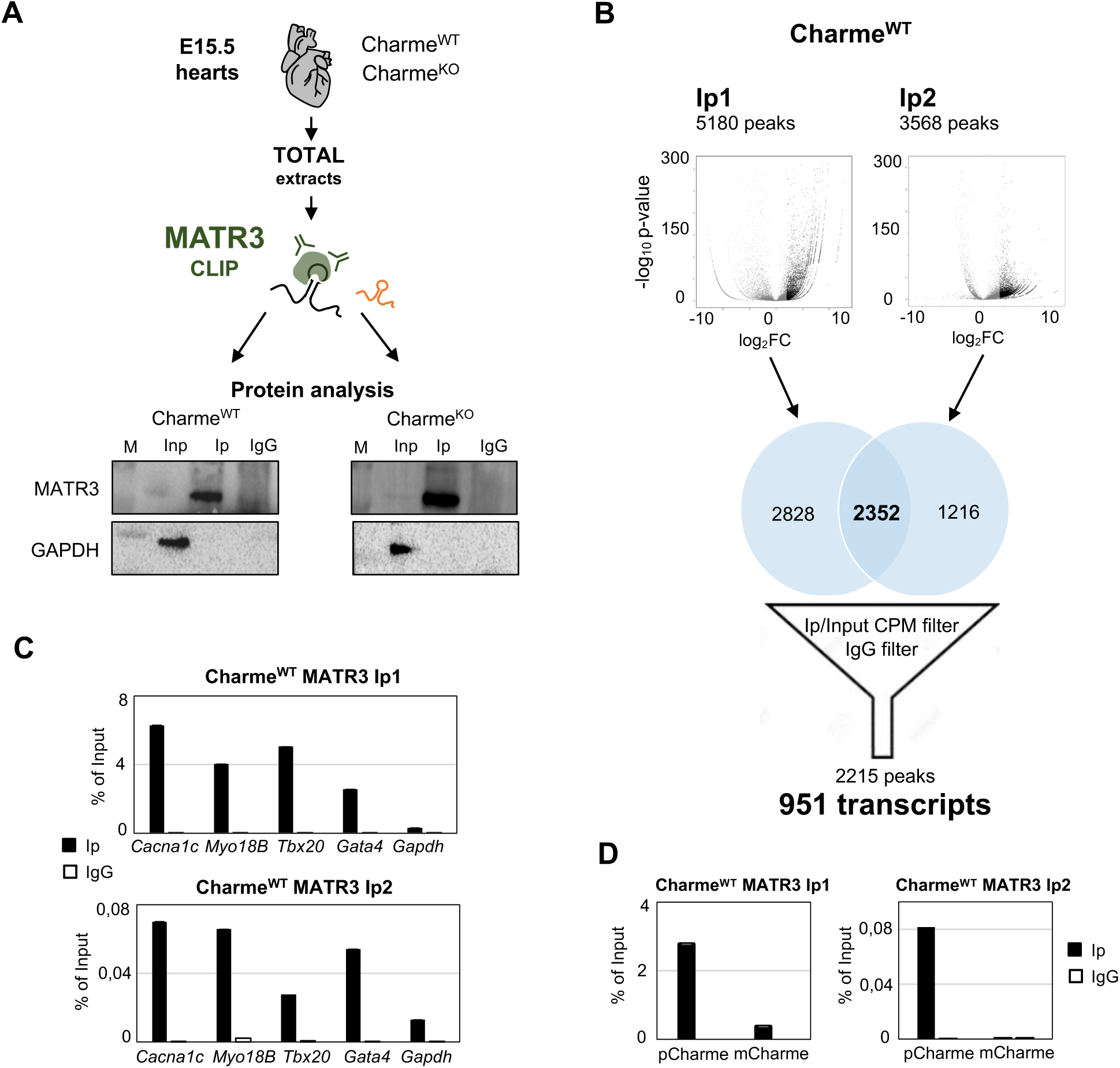
Related to Figure 4. **A)** Schematic representation of MATR3-CLIP assay as performed from fetal hearts (E15.5) in Charme^WT^ and Charme^KO^ conditions. MATR3 western blot analysis on the retrieved protein fractions is shown. GAPDH protein serves as a loading control. Input (Inp) samples represent 10% of the total protein extracts. **B)** Schematic representation of the workflow used for MATR3 CLIP-seq analysis. Volcano plots represent the fold-enrichment over Input (log_2_ Fold enrichment, x-axis) and significance (-log_10_ p-value, y-axis) of MATR3 peaks in the Ip1 (left panel) and Ip2 (right panel) samples. Black dots represent the significantly enriched peaks (log_2_ Fold enrichment > 2 and FDR < 0.05). Venn diagrams depict the intersection between Ip1 and Ip2 significantly enriched peaks. The 2215 filtered peaks correspond to the 951 MATR3-bound transcripts. CPM: Counts Per Million. See Materials and Methods for details. **C)** RT-qPCR quantification of *Cacna1c*, *Myo18B*, Tbx20 and *Gata4* RNA recovery in MATR3 Ip1 (upper panel), Ip2 (lower panel) and IgG Charme^WT^ samples. *Gapdh* transcript serves as negative control. Values are expressed as percentage (%) of Input. **D)** RT-qPCR quantification of pCharme and mCharme RNA recovery in MATR3 Ip1 (left panel), Ip2 (right panel) and IgG Charme^WT^ samples. Values are expressed as percentage (%) of Input.

**Figure 5-Figure supplement 1.**
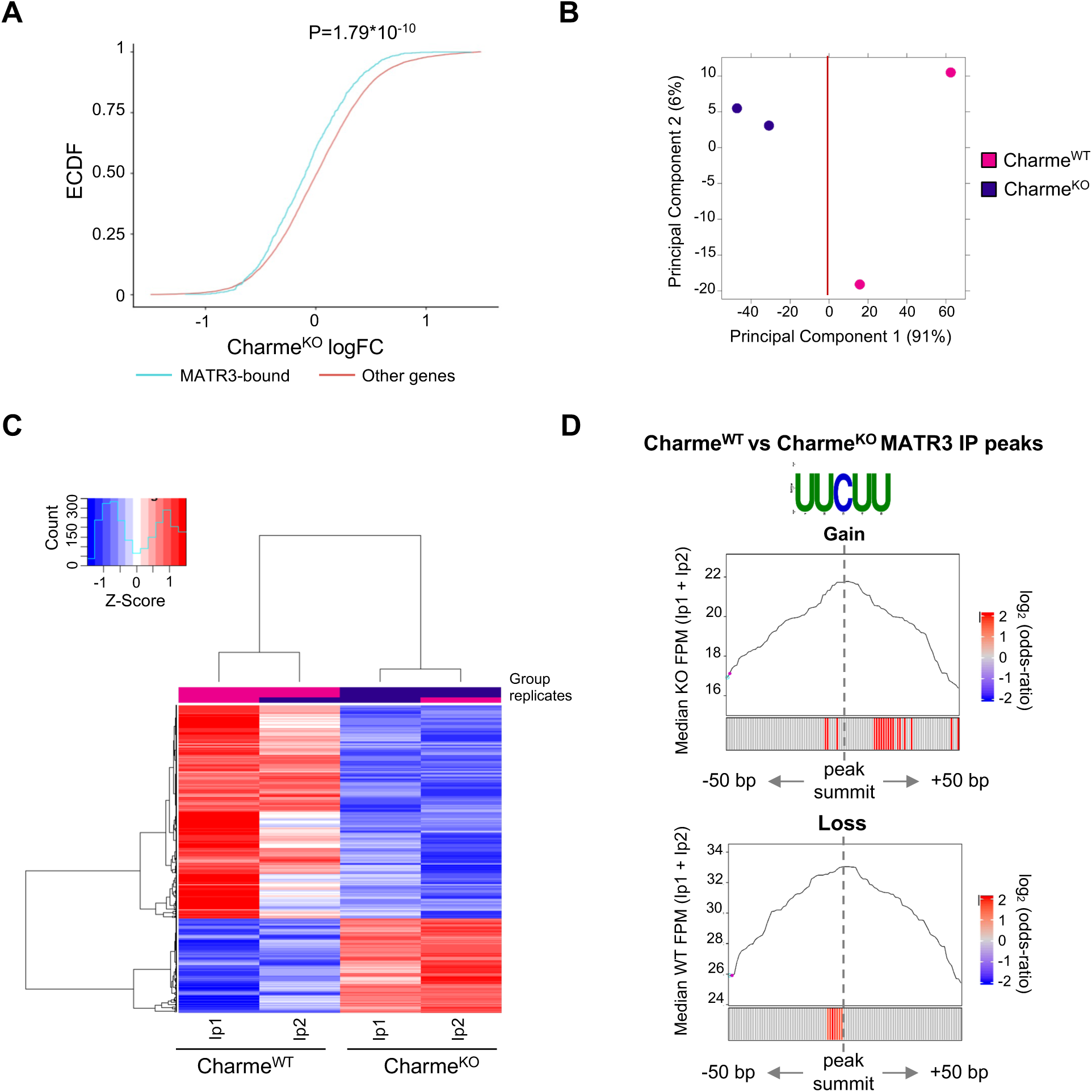
Related to Figure 5. **A)** Empirical Cumulative Distribution Functions (ECDF) showing the RNA abundance of MATR3 targets in Charme^KO^ condition compared to the other expressed genes. Significance was determined using a two-sided Kolmogorov–Smirnov (KS) test. **B)** Principal Component analysis (PCA) plot of Charme^WT^ and Charme^KO^ MATR3 CLIP-seq peaks. Plot was obtained from DiffBind package. See Material and Methods for details. **C)** Heatmap visualization of Charme^WT^ and Charme^KO^ MATR3 CLIP-seq peaks. Plot was produced by DiffBind package. Z-Score of normalized reads counts is shown. See Material and Methods for details. **D)** Positional enrichment analysis of MATR3 motif in differential MATR3 CLIP-seq peaks (Charme^WT^ *vs* Charme^KO^). For each analysed position close to peak summit, line plot displays the median CLIP-seq signal (FPM, IP1 + IP2) while heatmap displays the log_2_ odds-ratio of UUCUU motif enrichment. Significant enrichments (p-value < 0.05) are shown in red. See Material and Methods for details. CLIP-seq signal in Charme*^KO^* and Charme^WT^ conditions is displayed for «Gain» and «Loss» peaks, respectively.

